# PhyloCNN: Improving tree representation and neural network architecture for deep learning from trees in phylodynamics and diversification studies

**DOI:** 10.1101/2024.12.13.628187

**Authors:** Manolo Fernandez Perez, Olivier Gascuel

## Abstract

Phylodynamics and diversification studies using complex evolutionary models can be challenging, especially with traditional likelihood-based approaches. As an alternative, likelihood-free simulation-based approaches have been proposed due to their ability to incorporate complex models and scenarios. Here, we propose a new simulation-based deep learning (DL) method capable of analyzing large datasets and accurately estimating parameter values for birth-death models in both phylodynamics and diversification studies. Our approach involves encoding trees by extracting a vector of local features for all nodes of the input phylogeny. We also developed a dedicated convolutional neural network architecture called PhyloCNN. Using simulations, we compared the accuracy of PhyloCNN when using feature vectors with a variable number of generations to describe the local context of nodes and leaves. The number of generations had a greater impact when considering smaller training sets, with a broader context showing higher accuracy, especially for complex evolutionary models. Compared to other recently developed DL approaches, PhyloCNN showed higher or similar accuracies for all parameters when used with training sets one or two orders of magnitude smaller (10,000 to 100,000 simulated training trees, instead of millions). We applied PhyloCNN with compelling results to two real-world phylodynamics and diversification datasets, related to HIV superspreaders in Zurich and to primates and their ecological role as seed dispersers. The high accuracy and computational efficiency of our method opens new possibilities for phylodynamics and diversification studies that need to account for idiosyncratic phylogenetic histories with specific parameter spaces and sampling scenarios not considered in more general approaches.

## Introduction

The shape of phylogenetic trees, characterised by their topology and branch lengths, can be used as a source of information about the underlying evolutionary processes responsible for generating these phylogenetic relationships. Phylodynamics and diversification dynamics studies combine this phylogenetic information with stochastic modelling to infer key parameters about pathogen transmission (Grenfell et al., 2004; Stadler, 2010; Volz et al., 2013; Featherstone et al., 2022) and species diversification (Maddison et al., 2007; Morlon et al., 2010; Morlon, 2014) over time. With the advent of affordable, high-throughput sequencing, there has been a massive increase in the number of individuals sampled and genomic regions sequenced in epidemiological (Attwood et al., 2022) and evolutionary (Garrick et al., 2015) studies. In phylodynamics, this makes it possible to obtain sequences on the same time scale as the transmission and to estimate epidemiological parameters during disease outbreaks. For example, Vaughan et al. (2024) analyzed the branching patterns of SARS-CoV-2 phylogenies to infer critical parameters such as the basic reproduction number (*R*_0_), the cumulative number of infections, and the effectiveness of public health interventions during the early outbreaks of the COVID-19 pandemic around the world. In macroevolution, Condamine et al. (2019) examined the diversification rate of 218 tetrapod families and showed the prevalence of diversification slowdowns caused by both climatic cooling and global diversity changes.

Beyond coalescent theory, models and predictions (Volz et al. 2013; Müller et al. 2017), understanding the mechanisms driving the dynamics of pathogen spread and species diversification has often relied on the development of birth-death models (Nee, 2006; Stadler, 2010). Such models provide a probabilistic framework to study pathogen transmission (or birth, speciation) and host recovery (or death, extinction) processes, including a number of factors, in particular sampling rate, size and structure of the population, and external drivers such as temperature and altitude (Höhna et al., 2011; Etienne et al., 2012; Pennell and Harmon, 2013; Stadler and Bonhoeffer 2013; Rabosky, 2014; Beaulieu and O’Meara, 2015; Sanmartín and Meseguer, 2016; Featherstone et al., 2023; Morlon et al., 2024). Given their similarities, MacPherson et al. (2022) have proposed a unifying framework for birth-death models in both epidemiology and macroevolution, establishing connections among their numerous variants. However, while these models have the ability to capture a wide range of questions in evolutionary biology, they pose many difficulties from a mathematical and computational point of view. In a recent paper, Louca and Pennell (2020a) discussed the issue of identifiability and showed that, in some models, certain parameters cannot be estimated from data without additional constraints and knowledge. Another major computational issue is that of likelihood calculation. For most models, the likelihood value of a given phylogeny and set of parameter values is the solution of a large recursive system of ordinary differential equations (ODEs), which is computationally expensive to solve and prone to numerical errors (Louca and Pennell, 2020b; Scire et al., 2022; Zhukova et al., 2023). Just as likelihood calculations can be problematic, so can parameter estimation in maximum-likelihood and Bayesian frameworks. In this context, the application of likelihood-based approaches is generally computationally intractable for most birth-death models and large amounts of data (DeMaio et al., 2015; Müller et al., 2017), such as those obtained by advanced omics technologies.

To circumvent these limitations, likelihood-free approaches based on simulations have been proposed. Within this suite of methods, Approximate Bayesian Computation (ABC) is the most frequently used. For ABC, the input data (i.e., a phylogenetic tree in our context) is first reduced to a set of summary statistics (SumStats), which are then used in a rejection step that compares the input SumStats to simulated ones (Beaumont et al., 2002; Csilléry et al., 2010). Although ABC is powerful, there are several drawbacks, such as the use of a rejection step that only retains a small proportion of simulations that are most similar to the empirical data (Csilléry et al., 2012). This is suboptimal, as the information from the remaining simulations is discarded. Moreover, the choice of the SumStats and the distance metrics (e.g., Euclidean) between simulated and real data can be rather arbitrary (Robert et al., 2011; Pudlo et al., 2016; Saulnier et al., 2017) and may fail to capture important aspects of the raw data, in addition to the risk of including issues related to the “curse of high dimensionality” (Prangle et al., 2018).

Some of these shortcomings are overcome by regression-based ABC approaches (Saulnier et al. 2017; Blum 2018). Following the same line, deep learning (DL) methods have started to be applied to population genetics (Sheehan and Song, 2016; Schrider and Kern, 2018), species delimitation (Perez et al., 2022), phylogenetics (Smith and Hahn, 2023; Mo et al., 2024), phylodynamics (Voznica et al., 2022), macroevolution (Lambert et al., 2023), and phylogeography (Kirschner et al. 2022; Thompson et al., 2024). Unlike standard ABC, they use “pattern-recognition” DL techniques to discover meaningful features from the raw data (Flagel et al., 2019; Torada et al., 2019). Moreover, since DL architectures are usually trained on a large number of simulations, it is possible to evaluate complex evolutionary scenarios, considering multiple parameters and mirroring the idiosyncrasies of the empirical dataset, such as temporal and spatial sampling bias (Danesh et al. 2023). Another striking advantage of DL is its flexibility to incorporate multiple data types (e.g., phenotypic and environmental traits, sequence alignments, phylogenies, global summary statistics).

Several DL applications in population genetics and phylogeography have used either SumStats (Mondal et al., 2019) or raw alignments of SNPs (Flagel et al., 2019; Oliveira et al., 2020; Sanchez et al., 2021). However, some recent studies suggest the usefulness of analysing phylogenetic information by directly encoding the tree shape and applying convolutional neural networks (CNN; Voznica et al., 2022; Lambert et al., 2023; Thompson et al. 2024). These methods are based on the Compact Bijective Ladderized Vector (CBLV) encoding of trees that we proposed in (Voznica et al. 2022). This combinatorial tree representation encodes a rooted tree with *n* leaves and 2*n* - 2 branch lengths into a numerical vector of length 2*n* - 1. This is a very compact tree representation because we encode not only the branch lengths but also the tree topology. However, when we designed this tree representation, we did not have a particular DL architecture in mind. While high accuracy was achieved, higher than state-of-the-art likelihood-based software (Voznica et al., 2022; Lambert et al., 2023), the performance was no better than using SumStats (when available, as in phylodynamics) and a standard feed-forward neural network (FFNN; Haykin, 1999). Moreover, training a new model required millions of simulated trees due to the complexity of the CBLV representation and its combination with CNN (hereafter CNN_CBLV), where important features are available because the representation of the tree is complete, but are somehow hidden and difficult to extract due to the combinatorial nature of the encoding. Therefore, while the implementation of any birth-death model using CBLV is conceptually simple since there is no need to design new SumStats, the computational cost is high. This limits the exploration of new ideas and models, as well as new data with specific characteristics that would require simulating millions of trees and (re)training the neural architecture on those trees.

Here we propose the combination of a new tree representation and a dedicated CNN architecture, applicable to both phylodynamics and diversification models. Both the tree representation and the architecture were designed jointly, resulting in higher accuracy (significantly better than FFNN using SumStats - hereafter FFNN_SumStats) with 10 to 100 thousand trees (instead of 4 million with CBLV in Voznica et al., 2022, and 1 million in Lambert et al., 2023). This opens the way for exploratory analyses of new models and a wide range of studies. In the following, we first introduce this tree representation and the corresponding CNN architecture, called PhyloCNN. We then evaluate the performance of PhyloCNN using simulated data and perform comparisons with state-of-the-art programs, PhyloDeep (Voznica et al., 2022) for phylodynamics and Lambert et al.’s (2023) approach for diversification (hereafter named DeepTimeLearning - DTL, as per its GitHub repository). To make comparisons more straightforward, we applied PhyloCNN to the same empirical datasets, using both a phylodynamic HIV dataset (men-having-sex-with-men in Zurich; Rasmussen et al., 2017, also used with PhyloDeep) and a diversification study (primates and their ecological role as seed dispersers; Gómez and Verdú, 2012, also used with DTL).

## Models and methods

Our work was focused on the development of an improved DL approach (PhyloCNN) for the analysis of phylogenetic trees under different birth-death models for phylodynamics and macroevolution studies. To this end, and to allow comparisons of PhyloCNN with previously developed DL approaches, we have structured this section according to the following outline: (1) description of the different birth-death models considered for both phylodynamics and macroevolution; (2) procedures for simulating phylogenetic trees under each birth-death model; (3) PhyloCNN tree representation; (4) neural network architectures for model selection and parameter estimation, training criteria and algorithms; (5) criteria used to compare PhyloCNN with other approaches; (6) application of PhyloCNN to empirical datasets.

### Birth-Death models for phylodynamics and macroevolution

A large number of birth-death models have been developed for both phylodynamics and diversification dynamics (see MacPherson et al., 2022, for an overview of the models and the connections among their variants). Since our goal was to develop a better tree representation along with a more appropriate DL architecture and compare it to state-of-the-art approaches, we focused on a subset of birth-death models used in (Voznica et al., 2022, phylodynamics) and (Lambert et al., 2023, diversification dynamics).

In phylodynamics, this corresponds to three well-established models: a simple birth-death model with incomplete sampling (BD; Fig. 1(a); Stadler, 2010), a birth-death model with an exposed class and a rate ε to become infectious (BDEI, Fig. 1(b); Stadler et al., 2014) and a birth-death model with superspreading (BDSS, Fig. 1(c); Stadler and Bonhoeffer, 2013). In these three models, γ is the removal rate, and if a patient is removed, it has a probability *s* of being sampled. In these three models, the β parameters correspond to transmission events, whose rate with BDSS depends on the spreading state (super-spreader = S, normal-spreader = N). In BDSS, we are interested in the fraction of super-spreaders (*f*_SS_) in the population and in the superspreading ratio (*X*_SS_), which compares the transmission rate of super-spreaders to the rate of normal-spreaders. As in (Voznica et al., 2022), we imposed constraints on the β parameter values with BDSS, corresponding to the fact that the spreading state of the recipient does not depend on the spreading state of the donor (Fig. 1(c)).

**Figure 1:**
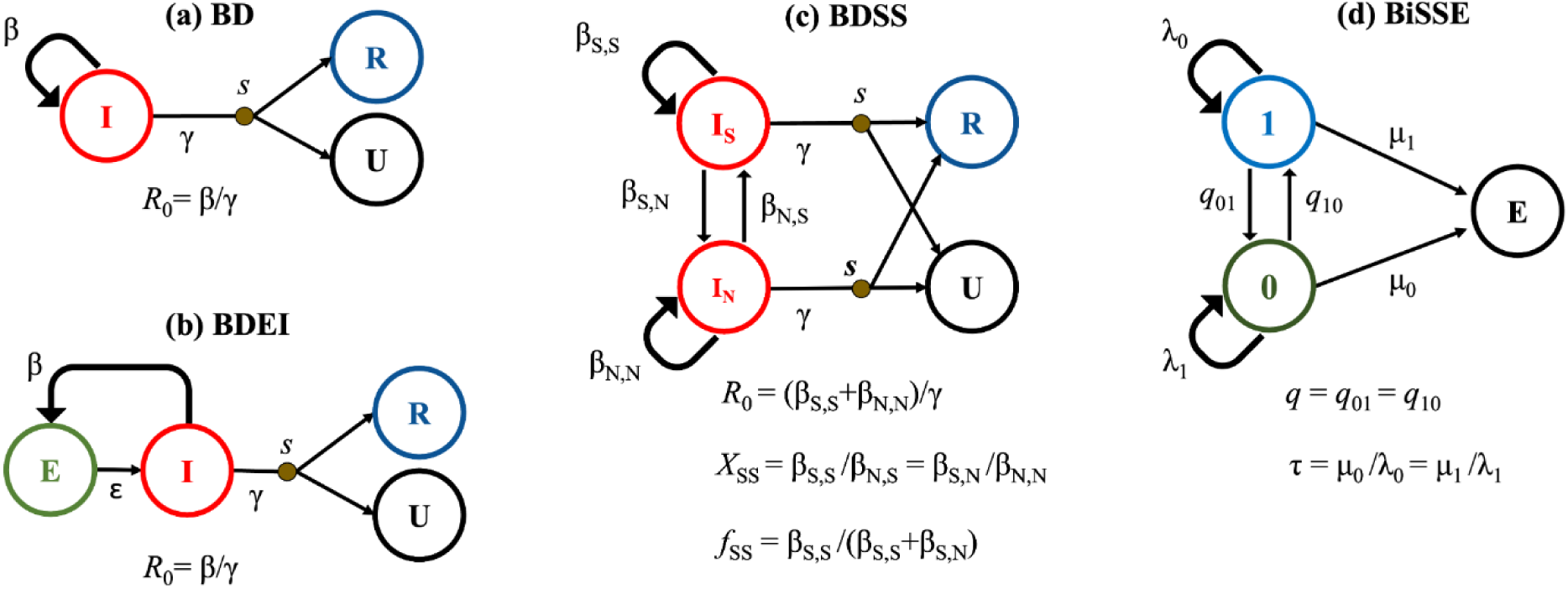
Birth-death models with incomplete sampling implemented in PhyloCNN. Phylodynamics: (a), (b), and (c); Macroevolution: (d). **(a)** The Birth Death (BD) model is the simplest model for phylodynamics and includes three classes of individuals: the infected (I) who transmit the disease at rate β and are removed at rate γ and then sampled (R) or left unsampled (U) with probability *s*. This model can be used to estimate *R*_0_ (= β/γ) and the infectious period (1/γ). **(b)** The BD model with exposed and infectious individuals (BDEI) includes the exposed class (E), which is associated with an incubation rate ε to become infectious and is used to estimate the additional latency period parameter (1/ε). **(c)** The superspreading model (BDSS) extends the BD model by separating the infected individuals into one class for super-spreaders (I_S_) and another for normal-spreaders (I_N_). The super-spreader individuals are a low subset of the infected population (with fraction *f*_SS_) that transmit the disease at a higher rate (*X_SS_*) than the normal-spreaders. There are specific transmission rates for each possible pair of donor-recipient (super-spreader or normal-spreader) individuals (i.e., β_S,S_, β_S,N_, β_N,S_ and β_N,N_), which are constrained by equality β_S,S_/β_S,N_ = β_N,S_/β_N,N_, which basically means that the recipient’s state does not depend on the donor’s state. **(d)** The BiSSE model is a state-dependent BD model for diversification studies, with two binary discrete classes (0 and 1) that generate new species with their same character state at rates λ_0_ and λ_1_, respectively. Class transitions occur with symmetric rates *q* = *q*_01_ = *q*_10_ along the tree branches (anagenetically) and species from each state become extinct (class E) with rates μ_0_ and μ_1_. The turnover rate (τ) and the sampling probability are shared among states. The outline of the models was adapted from (Voznica et al., 2022).

The binary state speciation and extinction (BiSSE) model (Fig. 1(d)) applies to diversification studies and is a simple state-dependent BD model that shares similarities with BDSS. As in BDSS, species are characterised by a binary state (0 or 1), which can affect their constant rates of speciation (λ_0_ and λ_1_) and extinction (μ_0_ and μ_1_). Species can transition anagenetically from one state to another along branches (at rates *q*_01_ and *q*_10_; unlike the BDSS, transitions do not occur during speciation/transmission events). Extinction events (μ_0_ and μ_1_) lead to the extinct class (E) of species not sampled and not visible in the observed tree. The sampling scheme follows a Bernoulli distribution at the present time (after the tree has grown) according to the sampling probability, allowing the analysis of trees in which some, but not all, of the extant species are present. This is an additional difference from phylodynamics, where the patients (tree leaves) are sampled continuously over time. As in (Lambert et al., 2023), we consider a simple version of BiSSE with symmetric transition rates (*q* = *q*_01_= *q*_10_) as well as turnover rate and sampling probability at present (τ and *s*, respectively) that are shared across species regardless of their state.

### Simulating datasets for phylodynamics and macroevolution

To train and test our network architectures, we simulated large sets of trees that were recorded along with the parameter values used to generate those trees. The phylodynamics datasets under the BD, BDEI and BDSS models were simulated using the scripts from (Voznica et al., 2022). For diversification dynamics, we simulated datasets under the BiSSE model using castor v1.6.6 (Louca and Doebeli, 2018) with the same script and options used in (Lambert et al., 2023). To simplify the comparison of PhyloCNN with these previous studies, we used the same tree sizes (200-500 tips) and priors for all parameters (Supp. Tab. S1 and S2, recovered from Supp. Tab. S4 in (Voznica et al., 2022) and Tab. 2 in (Lambert et al., 2023)). These priors are uniform and non-informative for the phylodynamics models. With BiSSE, the simulation procedure used in (Lambert et al., 2023), which we also used here, ensures that state 0 always corresponds to the most frequent state. It follows that λ_0_ is generally (∼80% of the trees) larger than λ_1_, and the priors are non-uniform and somehow informative (the λ_0_ and τ priors are uniform, but λ_1_’s and *q*’s are not; Fig.7, Supp. Tab. S2).

We simulated 100 thousand (K) trees per model. These trees were used to train PhyloCNN to perform parameter estimation under each model considered, using a split for training and validation sets of 70% and 30%, respectively. We also performed experiments using only 10K trees (extracted from the 100K trees) to train PhyloCNN to estimate parameters, using the same split percentages for training and validation. To perform model selection (classification) for BD, BDEI and BDSS in phylodynamics, we trained PhyloCNN with 10K trees per model (also using the same split percentages for training and validation). We evaluated the different methods for phylodynamics on a set of 1K test trees extracted from the 10K trees for each model used as the test set for PhyloDeep in (Voznica et al., 2022). Since the test trees from DeepTimeLearning were not provided, to test PhyloCNN for diversification inference, we simulated 1K trees using castor and the same options and priors as for the training trees.

### Encoding phylogenies based on node neighborhood

Here, we propose a new approach to encoding phylogenies that extracts meaningful information and measurements from the neighborhood of each node and can be used to train deep learning algorithms to perform model selection (classification) and parameter estimation (regression) from both phylodynamics and macroevolution datasets. To make our encoding independent of time scale, the trees (and model parameters) are rescaled to an average branch length equal to 1.0 (Voznica et al., 2022). PhyloCNN is trained on these rescaled trees and parameter values, which are rescaled back to the original time scale after prediction. The same procedure applies to the real input tree in the prediction phase.

Our encoding approach involves describing the neighborhood (e.g., length of outgoing branches) and main measurements (e.g., date, number of descendants) of all nodes and leaves of the phylogeny. Specifically, we compared the information extracted from three hierarchical levels of generational context by visiting each node in the tree (e.g., N in Fig. 2), with the python package ete3 v.3.1.3 (Huerta-Cepas et al., 2016). These generational contexts include redundant information (e.g., the encoding of node N includes information on other nodes that will also be encoded, such as its ancestor A1 and children C1 and C2; Fig. 2). In this way, our encoding functions as a “windowing” approach, similar to methods used in computer vision where overlapping filters slide through the image. This windowing, often combined with CNN in computer vision, underlies our architectural choices (see below). As with CBLV, the representation is complete in the sense that the tree can be unambiguously reconstructed from its encoding, unless there are equalities of, for example, branch lengths, dates, and number of descendants, which are extremely rare in practice.

**Figure 2.**
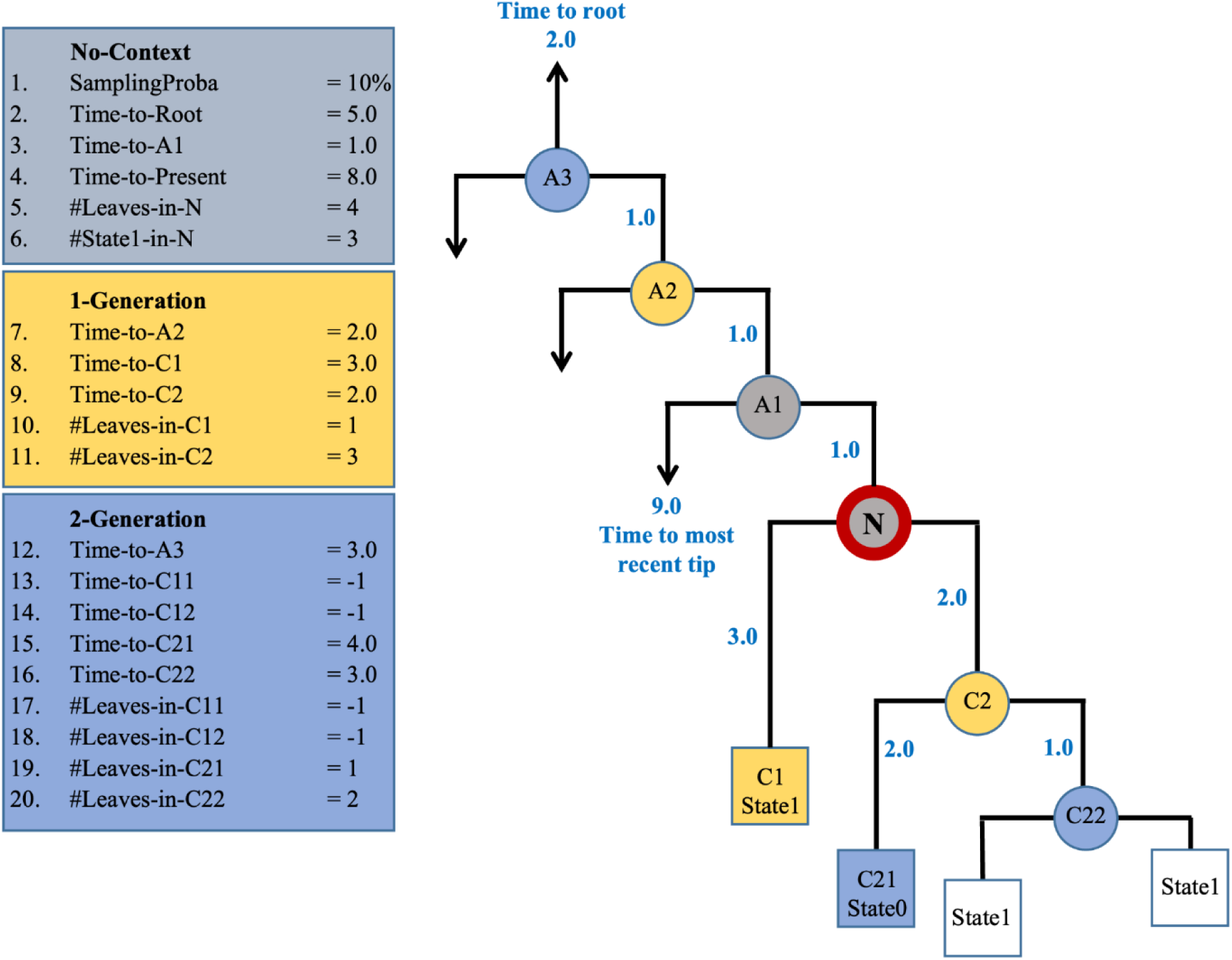
Encoding phylogenies with node neighborhoods. All internal nodes and leaves of the phylogeny are encoded by the procedure shown for node N (in red). In the first hierarchical level (No-Context; in grey), we encode the sampling probability for the phylogeny, the total number of leaves associated with N, the times to the root, to the parent node (A1) and to the present (the most recent tip in the tree). We also include the number of leaves in state 1 when considering the BiSSE model. In the next hierarchical level (1-Generation; in yellow), we record the times to the grandfather (A2) and child nodes (C1 and C2), along with the number of leaves associated with the latter. For the last hierarchical level (2-Generation context; in blue), we encode the time to the great-grandfather (A3) and the grandchildren (C11, C12, C21 and C22), together with the number of leaves associated with them. Missing values (e.g., measurements associated with N’s grandchildren “C11 and C12”, which are missing because C1 is a tip) are encoded as −1. Branch lengths are displayed in blue. The time to the most recent tip is shown as associated with A1 (9 time units in the example), which is used to calculate Time-to-Present for node N.

In the first hierarchical level (No-Context), we encode (Fig. 2):

1. The sampling probability of the whole phylogeny (SamplingProba, given as highly redundant information, repeated at all nodes).
2. The time from the node to the root (Time-to-Root).
3. The time to the parent (Time-to-A1; the branch length).
4. The time to the most recent tip in the tree (Time-to-Present).
5. The number of leaves descending from the node (#Leaves-in-N).
6. The number of descendant leaves in state 1 (#State1-in-N; only used with BiSSE; with leaves, this is equivalent to their trait state).

In the second hierarchical level (1-Generation, Fig. 2), we combine this information with the time (7) to the grandfather node (A2), the time (8) to the child (C1) with the lower number of descendant leaves, the time (9) to the child (C2) with the higher number of descendant leaves, the number (10) of leaves descending from C1 and the number (11) of leaves descending from C2. In the third hierarchical level (2-Generation, Fig. 2), we add the same information for the great grandfather node (A3) and N’s grandchildren (C11, C12, C21, and C22). These two higher-generation contexts (1- and 2-Generation) add a lot of information and in fact perform better than No-Context (see below), but also result in a notable number of unknowns (encoded with −1), which is likely to limit their performance on small trees where nodes have on average a limited number of ascendants and descendants (e.g. see Fig. 2, where using 2-Generation for node N induces 4 unknowns).

This information is encoded in a table, with each internal node/leaf in a row and each column containing one of the neighborhood features described above. After encoding, we import the table to Python v.3.9.18 and use NumPy v1.26.4 to split the information from internal nodes and leaves into two different tables and sort the rows (internal nodes and leaves) in ascending order of time (i.e., the first entry is the oldest internal node/leaf and the last entry is the most recent internal node/leaf). This standardization procedure was adopted because we observed slightly higher accuracy and faster convergence during training experiments. Since the total number of leaves exceeds the number of internal nodes by 1, we repeat the last entry for the internal nodes. This allows us to obtain two tables with the same dimensions for each tree. Also, since our approach is designed to analyze trees of different sizes, we pad the tables of leaves and internal nodes with zeros until we reach 500 rows (i.e., the number of leaves in our larger trees). This step is necessary because the CNN deep learning architecture requires that the input tables have the same size.

### Deep Neural Architectures and Training Algorithms

We developed a new neural network architecture to extract information from our proposed tree representation in Keras-Tensorflow v2.15.0. We use a depthwise separable convolutions architecture (Chollet, 2017), which consists of a depthwise 2D convolution layer with 32 neurons, and a kernel size corresponding to a (leaf, internal node) pair and the total number of encoded features (Fig. 2, 3). For this layer, we split the input into two channels: one for the table of encoded leaves, and the other for the internal nodes. We use the ‘groups=2’ option, which results in separate convolutions for each channel. This procedure was inspired by the fact that internal nodes and leaves contribute differently to the tree likelihood calculation for multi-type birth-death models (MTBD, which includes BD, BDEI and BDSS; see Equation 8 in Zhukova et al., 2023). This layer is followed by a pointwise layer that uses 32 1 × 1 filters to combine the outputs from the two different channels. This is followed by an additional non-linear convolution layer with 32 1 × 1 filters, as recommended in (Chollet, 2017). The resulting feature maps are flattened (collapsed into a single linear vector) with a 2D global average pooling layer. The resulting vector is then fed into a FFNN with 4 fully connected (FC) layers with decreasing numbers of neurons (64, 32, 16, 8) and an additional output FC layer with a number of output neurons equal to the number of parameters to be estimated (4 for BiSSE and BDSS, 3 for BDEI, and 2 for BD). For model selection (classification), the only modification to the architecture is the output layer, for which we use a softmax layer with 3 neurons corresponding to the three phylodynamics models.

**Figure 3.**
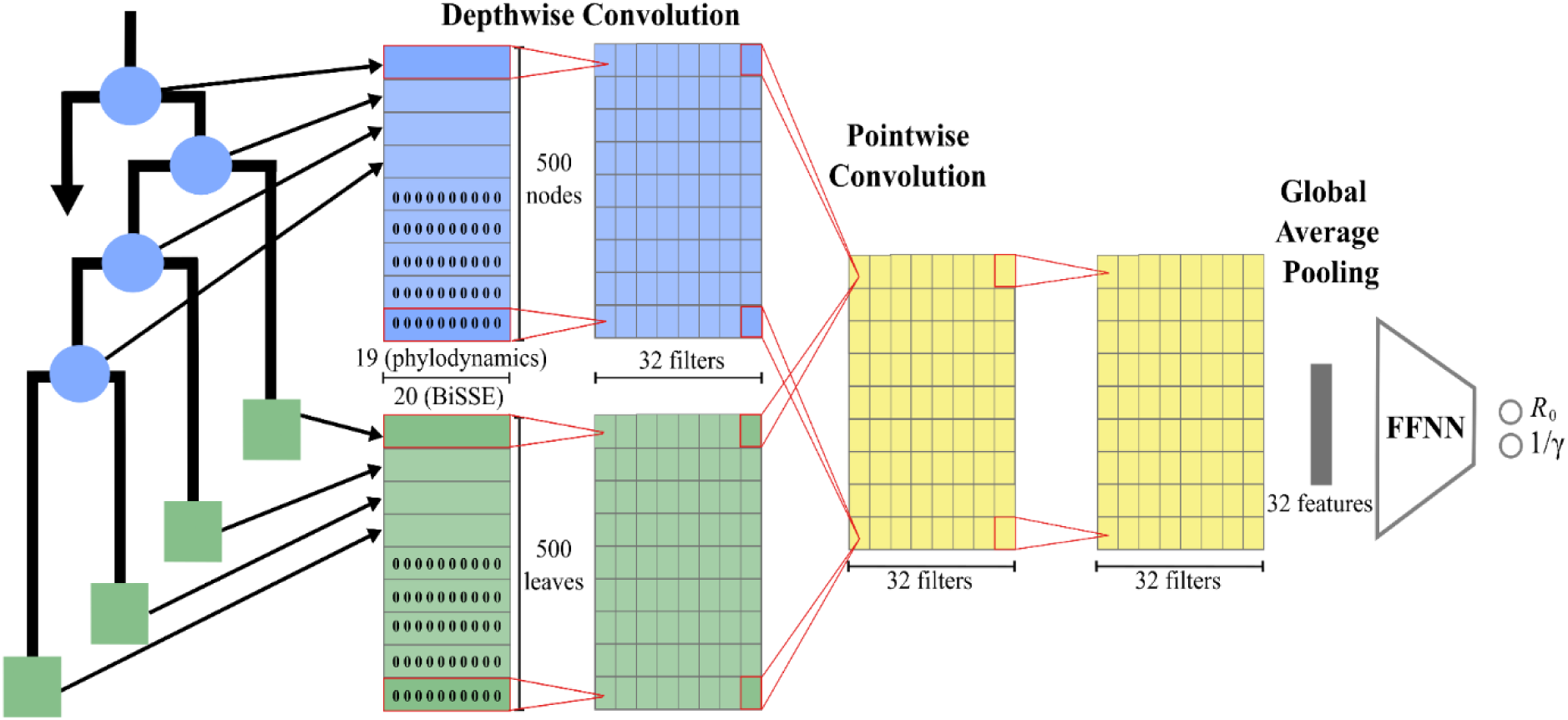
PhyloCNN architecture. The encoding tables for the internal nodes and the leaves of the phylogeny are entered in separate channels. Each contains a number of columns equal to the number of coding features and 500 rows (for trees with less than 500 leaves, the remaining rows are padded with 0s). The first layer performs depthwise convolutions (in each channel, with 32 filters of dimensions 1 × number of columns/features), followed by a pointwise convolution layer combining the two channels (with 32 1 × 1 filters) and an additional non-linear convolution layer (32 1 × 1 filters). The resulting 32 feature maps are flattened with global average pooling, giving in a single value for each of the 32 filter maps. The resulting vector is connected to an FFNN (with 64, 32, 16 and 8 neurons). Then follows the output layer with a number of neurons equal to the number of parameters to be estimated (4 for BiSSE and BDSS, 3 for BDEI and 2 for BD), or a softmax layer with 3 neurons (one for each phylodynamics model) for model selection (classification).

It is important to note that this architecture contains a relatively small number of weights, maintaining almost the same number of weights regardless of the complexity of the model and the number of input features from different generational contexts. Specifically, the number of weights ranges from 7,178 (for the BD model and No-Context) to 7,964 (for the BiSSE model with 2-Generation context). We refined the PhyloCNN architecture by starting with the one used in PhyloDeep and performing several fine-tuning runs with different architectures and hyperparameters, based on independent validation sets (not used for testing). The main gain was (consistently) obtained by using different channels for the internal nodes and leaves.

We used exponential linear units (ELUs) as nonlinear activation functions (Clevert et al., 2015) in all convolution and FFNN layers. The network parameters were optimized using the Adam optimizer (Kingma and Ba, 2014). The model was run for 1,000 epochs with early stopping and a patience of 100 epochs to prevent overfitting. The epoch with the highest validation accuracy was saved by model checkpointing. The training procedure was evaluated with the validation set using different loss functions, depending on the model being predicted, to facilitate comparison of PhyloCNN with previous studies. We used the mean absolute percentage error (MAPE; as in PhyloDeep) for parameter estimation under the phylodynamics models, the mean absolute error (MAE; as in DeepTimeLearning) for BiSSE, and the categorical cross-entropy loss function for model classification/selection.

### Methods comparison

We evaluated the accuracy of PhyloCNN with its three generational contexts to perform model selection by predicting 1K simulated test trees (not used for training or fine-tuning of the PhyloCNN architecture) for each phylodynamics model (BD, BDEI, BDSS). For each test tree, we compared the label associated with the generative model with the prediction obtained by PhyloCNN. We used the results to build a confusion matrix, with accuracy calculated as the probability of selecting the model from which the input data was simulated. This accuracy was compared to that reported in (Voznica et al., 2022) for PhyloDeep and BEAST2 (Bouckaert et al., 2014).

We compared the accuracy of PhyloCNN and its three generational contexts with that of PhyloDeep and DeepTimeLearning (DTL) to perform parameter estimation under each birth-death model. To do this, we evaluated the effect of varying the number of training trees used for each DL approach, since in machine learning theory (Haykin, 1999), accuracy (as well as, of course, running time and memory requirements) strongly depends on the sampling size. Specifically, we compared the pre-trained networks of PhyloDeep (based on 4 million (M) training trees) and DTL (1M training trees), with PhyloCNN trained on 100K trees or a subset of 10K trees. We also evaluated the accuracy of PhyloDeep retrained on the same 100K and 10K trees used to train PhyloCNN, as well as the accuracy of DTL retrained on the same 100K and 10K trees as PhyloCNN. For phylodynamics models, we evaluated both the CNN_CBLV and FFNN_SumStats versions of PhyloDeep. We used the same summary statistics as in the original publication, namely the 83 SumStats proposed by Saulnier et al. (2017), plus 14 on transmission chains and 1 on tree size. FFNN_SumStats showed the highest accuracy of all methods evaluated in the PhyloDeep study (Voznica et al., 2022), including BEAST2 (Bouckaert et al., 2014). For BiSSE, we also performed maximum-likelihood estimation (MLE) using the approach implemented in the R package diversitree 0.9-3 (Fitzjohn, 2012) on our newly generated 1K test trees. As in (Lambert et al., 2023), we did not use SumStats for BiSSE because the existing statistics for diversification studies (Janzen and Etienne, 2024) do not deal with trait values associated with tree leaves as used in BiSSE.

We evaluated the accuracy of each method using the test sets described above, which contain 1K simulations for each phylodynamics model (taken from Voznica et al., 2022) and 1K simulations for BiSSE (newly generated; see above for details). To facilitate comparisons, we used the same error measures proposed by Voznica et al. (2022) for phylodynamics, and by Lambert et al. (2023) for BiSSE. Specifically, for the phylodynamics parameters, we calculated the mean relative error (MRE):

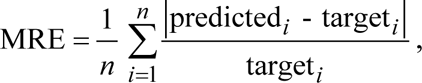

where *i* represents one test tree and *n* is the number of test trees (i.e., 1,000). We also calculated the mean relative bias (MRB) for all models and parameters:

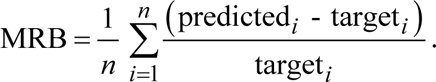

In the case of BiSSE parameters, we calculated the mean absolute error (MAE) and the mean bias (MB), as in (Lambert et al., 2023):

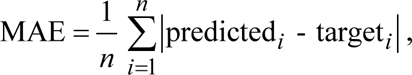

and

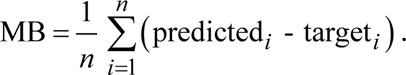

### HIV phylodynamics dataset

For direct comparison, we used the same tree evaluated in (Voznica et al., 2022). This phylogenetic tree was reconstructed by Rasmussen et al. (2017) from 200 HIV-1 sequences associated with the largest cluster of HIV-infected men who have sex with men (MSM) dataset, collected as part of the Swiss Cohort Study (2010). These data provide a snapshot of HIV-1 transmission within a highly interconnected subpopulation, allowing the phylodynamics of the virus in a local sexual contact network to be studied. When analyzed with PhyloDeep, the BDSS model was selected as the scenario that better fits the data (Voznica et al., 2022). Under this model, the posterior distribution of the infectious period (1/γ) using the CNN_CBLV approach showed values (95% CI = [8.1, 12.3]) outside the prior bounds (uniform from 1.0 to 10.0) used to train PhyloDeep. In addition, the posterior obtained for the fraction of super-spreaders (*X*_SS_) showed a distribution shifted toward the highest limit of the prior (95% CI = [6.7, 10.0]), with uniform prior bounds from 3.0 to 10.0, and a much higher value when analyzed with BEAST (95% HPD = [8.0, 26.1], Voznica et al., 2022).

For this reason, in the simulations we used custom prior distributions for 1/γ and *X*_SS_ by increasing their upper bounds (15.0 and 30.0, respectively; see Supp. Tab. S1 for details). We also used a uniform prior ranging from 0.2 to 0.3 (as suggested by Voznica et al., 2022) for the sampling probability, and we fixed the tree size to be the same as that of the HIV dataset (i.e., 200 leaves). We performed model selection on this HIV dataset using a training set of 3 x 10K trees simulated under each phylodynamics model and the new priors. We also performed an *a priori* check of model adequacy using the 10K trees simulated under the new priors and the BDSS model, which was consistently selected by PhyloCNN. First, we performed a principal component analysis (PCA) on the SumStats of both simulations and the HIV phylogeny (Supp. Fig. S2). We also evaluated each of the SumStats by checking whether the corresponding value for the HIV phylogeny was between the minimum and maximum values observed in the training set. Then, we simulated a larger training set of 50K trees of the same size under the selected BDSS model and the new priors to perform parameter estimation. Based on the simulation results (see below), 50K training trees seems to be a good compromise between computational efficiency and accuracy. In addition, 50K training trees makes it possible to draw reliable posteriors for each of the parameters and go beyond simple point estimates. We also compared PhyloCNN with FFNN_SumStats trained on the same set of 50K trees. Performing the same experiment with new priors and fixed tree size would be hardly manageable with CNN_CBLV, which typically requires millions of training trees (Voznica et al., 2022).

Finally, we performed an *a posteriori* model adequacy check using an approach similar to that of (Voznica et al., 2022). We simulated an independent set of 10K trees under the BDSS model, sampling parameter values from their posterior distributions obtained by PhyloCNN in the HIV tree analysis. We then performed a PCA on the SumStats of the simulated trees thus obtained and those of the HIV tree (Supp. Fig. S3). We also compared, for each SumStat, its value for the HIV tree and the minimum and maximum values obtained in these *a posteriori* simulations (see Supp. Material and Methods for details).

### Primates diversification dataset

We also ran PhyloCNN on an empirical diversification dynamics dataset. Again, to ensure a direct comparison with DeepTimeLearning, we used the same phylogenetic tree as in (Lambert et al., 2023). This tree consists of a subtree of the primates phylogeny from (Fabre et al., 2009) associated with trait information from (Gómez and Verdú, 2012) on the 260 species studied. Specifically, primate species were classified according to their mutualistic (fruit consumption and seed dispersal) or antagonistic (destructive) interactions with plants, corresponding to states 0 and 1 in the BiSSE model, respectively. For this dataset, we used the version of PhyloCNN trained under BiSSE, with the same priors as described above for the simulated data. The sampling fraction of the primates tree was set to 0.68, following the procedure described in (Lambert et al., 2023). We performed the same *a posteriori* model adequacy check described above for the HIV dataset (Supp. Fig. S4).

## Results

### Runtime and storage requirement

Details on the runtime and storage throughout our various experiments are available in the Supplementary Material (Tab. S1). Here, we present the main results and comparisons.

To simulate 100K trees per model for training PhyloCNN, we needed a runtime of ∼13 CPU hours for phylodynamics models (compared to ∼800 CPU hours for the 4M trees used in PhyloDeep; Voznica et al., 2022) and ∼4 CPU hours for BiSSE (compared to ∼40 CPU hours for the 1M trees used in DeepTimeLearning (DTL); Lambert et al., 2023). PhyloCNN tree encoding requires slightly more runtime per tree (∼2e-5 CPU hours versus ∼1.5e-5 CPU hours for CBLV), but the total time required to encode the PhyloCNN training set (100K trees per model; ∼2 CPU hours) is still much less than the time required for PhyloDeep (4M trees per model; ∼60 CPU hours) and DTL (1M trees; ∼15 CPU hours).

Another important point to consider is the storage required for the simulated trees (∼500 MB and ∼700 MB for 100K trees per phylodynamics and BiSSE model, respectively). Since PhyloCNN uses at least ten times fewer trees for training, the required storage is much smaller than the alternative DL approaches. However, the storage requirement for the encoded PhyloCNN trees is much higher, especially when using the 2-Generation context (∼10 GB for 100K trees per model), because our encoding is highly redundant, with 9 branches described for each tree node. CBLV is much more compact, as each node is described only once, resulting in ten times less storage per tree (∼1 GB for 100K trees). However, such a difference is compensated by the smaller number (at least ten times less) of trees encoded in the training set of PhyloCNN. Moreover, trees can be encoded on the fly during the learning phase.

Training PhyloCNN on 100K trees also required shorter runtimes (∼2 hours on Nvidia Tesla V100 GPUs) than PhyloDeep on 4M trees (∼5 hours for FFNN_SumStats and ∼150 hours for CNN_CBLV under BDSS on Nvidia Titan X GPUs; Voznica et al., 2022) and DTL on 1M trees (∼8 hours on Nvidia Titan X GPUs; Lambert et al., 2023). Although these GPUs are of a different generation (Nvdia Tesla V100 is ∼60% faster than Nvidia Titan X), the difference of PhyloCNN with CNN_CBLV is very significant, probably due to the combinatorial nature of this representation and the difficulty of extracting relevant features.

The runtime required to simulate 50K BDSS trees for the HIV study, encode them, and train PhyloCNN was ∼11 hours (∼6.5 for simulation, 1 for encoding, and 3.5 for training) on a MacBook Pro (Apple M3 Pro with 11 core CPU, 14 core GPU and 16 core Neural Engine), compared to ∼80 hours of average runtime for BEAST2 with BDSS (Voznica et al., 2022), demonstrating the potential of the method to adapt to new datasets, models and priors.

### Performance on simulated phylodynamics datasets

We compared the performance of PhyloCNN and its three generational contexts (No-Context, 1-Generation, and 2-Generation) with the two versions of PhyloDeep using summary statistics (FFNN_SumStats) and the CBLV tree representation (CNN_CBLV). The results for the three phylodynamics models (BD, BDEI, and BDSS) are shown in Figure 4 and Supplementary Material (Tab. S4, S5, and Fig. S1).

**Figure 4:**
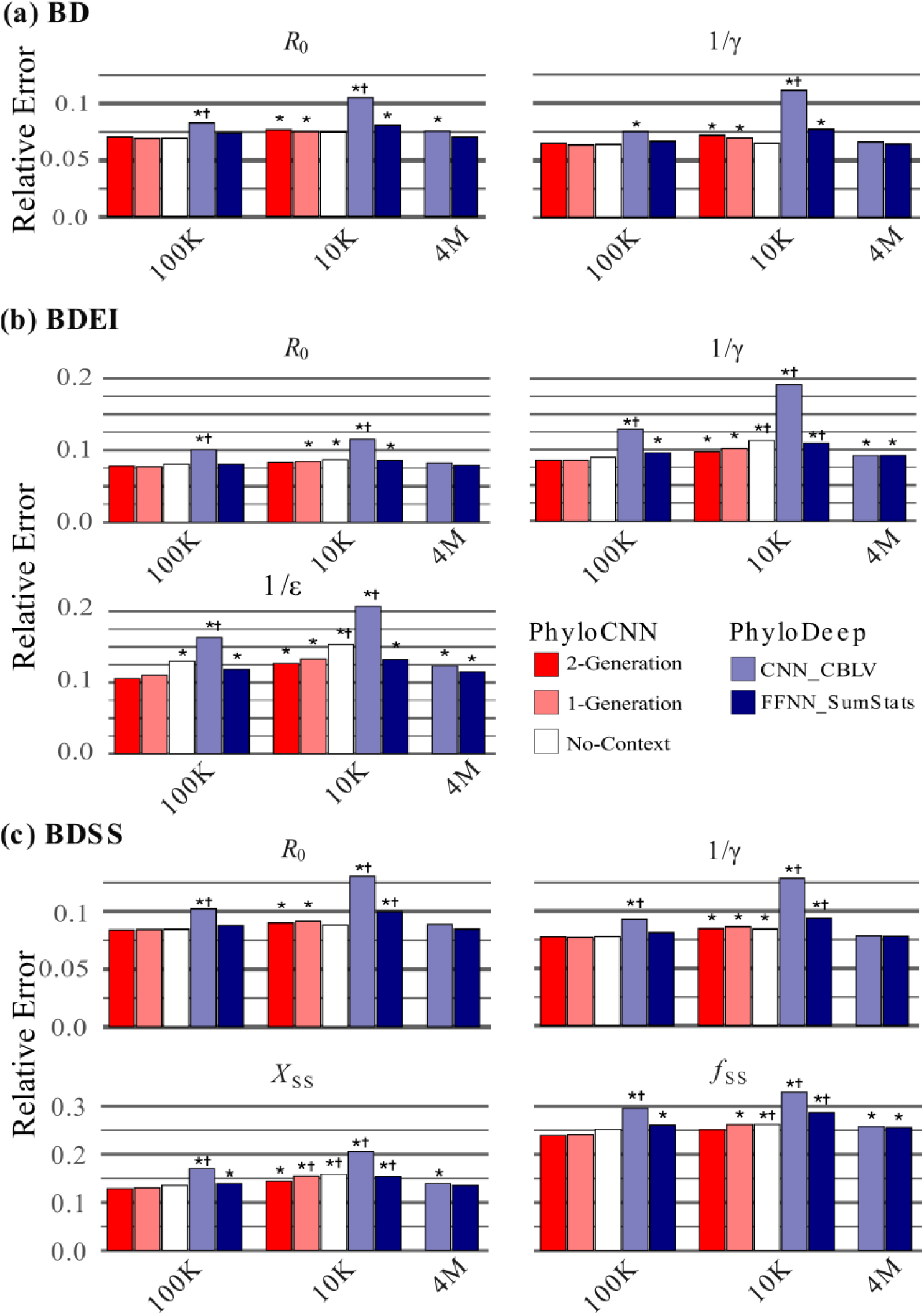
Mean relative error of PhyloCNN and PhyloDeep with phylodynamics models. Mean relative error (MRE) bars are shown for PhyloCNN with 2-Generation, 1-Generation and No-Context, using 100K and 10K training trees, and for the two PhyloDeep versions, CNN_CBLV and FFNN_SumStats, using 4M, 100K and 10K training trees. Throughout, we used the same 1,000 independent test trees for all methods. **(a)** BD model with parameters *R*_0_ and infectious period 1/γ. **(b)** BDEI model with the same two parameters and latency period 1/ε. **(c)** BDSS model with the two parameters estimated in BD together with the ratio of transmission rates between super- and normal-spreaders (*X*_SS_) and the fraction of super-spreaders (*f*_SS_). * Significantly higher MRE compared to PhyloCNN with 2-Generation and 100K training trees (paired Z-test, p<0.01). † Significantly higher MRE compared to PhyloCNN with 2-Generation and 10K training trees (p<0.01).

Model selection with PhyloCNN (Fig. S1) showed higher accuracy for both 1-Generation and 2-Generation than for No-Context (99.2%, 99.4%, and 89.6% correct predictions, respectively). Furthermore, both versions of PhyloCNN with context (trained on 10K trees per model) were significantly more accurate than PhyloDeep (trained on 4M trees per model) using FFNN with SumStats and CNN with CBLV (91.4% and 90.9%, respectively; Voznica et al., 2022).

Parameter estimation (Fig. 4) was more accurate for all DL methods when larger training sets were used, as expected from learning theory (Haykin, 1999). However, the effect is relatively small for FFNN_SumStats, which uses a simpler neural architecture and does not need to extract features from the raw data. The same is observed for the three generational contexts of PhyloCNN, which show a similar (relatively small) loss as FFNN_SumStats when trained with 10K trees instead of 100K, probably due to their relatively small number of neural network weights (∼7,000 to ∼8,000 versus ∼9,000 for FFNN_SumStats). On the contrary, and in agreement with Voznica et al. (2022), CNN_CBLV clearly needs very large training sets to be reasonably accurate (see, e.g., the results with 10K training trees, even with the simplest BD model; Fig. 4).

Among the three options of PhyloCNN, No-Context has the worst accuracy compared to 1-Generation and 2-Generation (Fig. 4, Sup. Tab. S4, S5). With 100K training trees, the differences are not significant (except for 1/ε with BDEI). With 10K training trees, the difference between No-Context and the other two becomes visible, especially for the most complex models (BDEI, BDSS) and the most difficult to predict parameters (1/ε, *f*_SS_, *X*_SS_). In these cases, No-Context seems to be handicapped by its very limited information on node features and neighborhoods, even though its CNN architecture contains (slightly) fewer weights to optimize than the other two options. The differences between 1-Generation and 2-Generation are small, especially with 100K training trees; 1-Generation seems to perform better with the simplest model (BD), but slightly worse with the two more complex ones (BDEI, BDSS).

PhyloCNN with 2-Generation consistently outperforms CNN_CBLV when trained with the same number of trees (100K or 10K; the differences in accuracy are significant for all models and all parameters, p<0.01). In fact, PhyloCNN with 2-Generation and 100K training trees has better accuracy than CNN_CBLV trained with 4M trees for all models and all parameters (5 among 9 significant differences, p<0.01). When PhyloCNN is trained with 10K trees, both become equivalent (no significant differences, p>0.01) for most parameters (except for 1/γ in both BD and BDSS for which PhyloCNN is significantly worse, p<0.01; Fig. 4, Sup. Tab. S4, S5).

PhyloCNN with 2-Generation also outperforms FFNN_SumStats when trained with the same number of trees (100K or 10K) for all models and all parameters, but to a lesser extent (differences in accuracy are significant for 9 out of 18, p<0.01). Compared to FFNN_SumStats trained with 4M trees, PhyloCNN with 2-Generation and 100K training trees is better or equal for all models and all parameters (3 significant differences among 9, p<0.01), while when trained with 10K trees only, it is no longer better, as expected considering the huge difference in training set size (4M versus 10K).

Overall, we observe relative errors of similar magnitude for the parameters *R*_0_ and 1/γ which are common to the three models, while the three model-specific parameters are more difficult to estimate, especially the BDSS parameters *X*_SS_ and *f*_SS_ (Fig. 4). These results are in line with the findings of (Voznica et al., 2022) and (Xie et al., 2024), which clearly indicate the difficulty of estimating the parameters of this complex model, where the distinction between normal and super-spreaders is hidden and not easily recoverable from the input phylogenetic tree alone. The results (Sup. Tab. S4 and S5) also show that all these DL methods have no tendency to induce a substantial bias for any of the models and parameters.

To summarize, PhyloCNN is significantly more accurate than both versions of PhyloDeep for the same number of training trees. It performs as well as or better than CNN_CBLV with 400 times fewer training trees (4M vs 10K) and FFNN_SumStats with 40 times fewer training trees (4M vs 100K).

### Performance on simulated diversification datasets

We compared the performance of PhyloCNN and its three generational contexts (No-Context, 1-Generation, and 2-Generation) with that of DeepTimeLearning (DTL). The later uses the PhyloDeep neural architecture and the CDV tree representation adapted from CBLV for diversification data such as those studied here in the setting of the BiSSE model (i.e., ultrametric trees with trait values attached to tree leaves; see also Thompson et al., 2024, for a similar adaptation in phylogeography). We did not run FFNN_SumStats because, to our knowledge, summary statistics available for diversification studies (e.g., Janzen and Etienne, 2024) do not deal with traits, or only marginally. For example, (Lambert et al., 2023) used adaptations of our phylodynamics summary statistics (Saulnier et al., 2017; Voznica et al., 2022) combined with four global summary statistics on traits (e.g., number of leaves in state 0 versus 1) to perform model adequacy checks, but not to predict parameter values. We also ran the diversitree software to perform maximum-likelihood estimation (MLE) of the parameters. The same 1K test trees generated under BiSSE were used for all methods. See above and Supplementary Material for details. The results are shown in Figure 5 and Supplementary Table S6.

**Figure 5.**
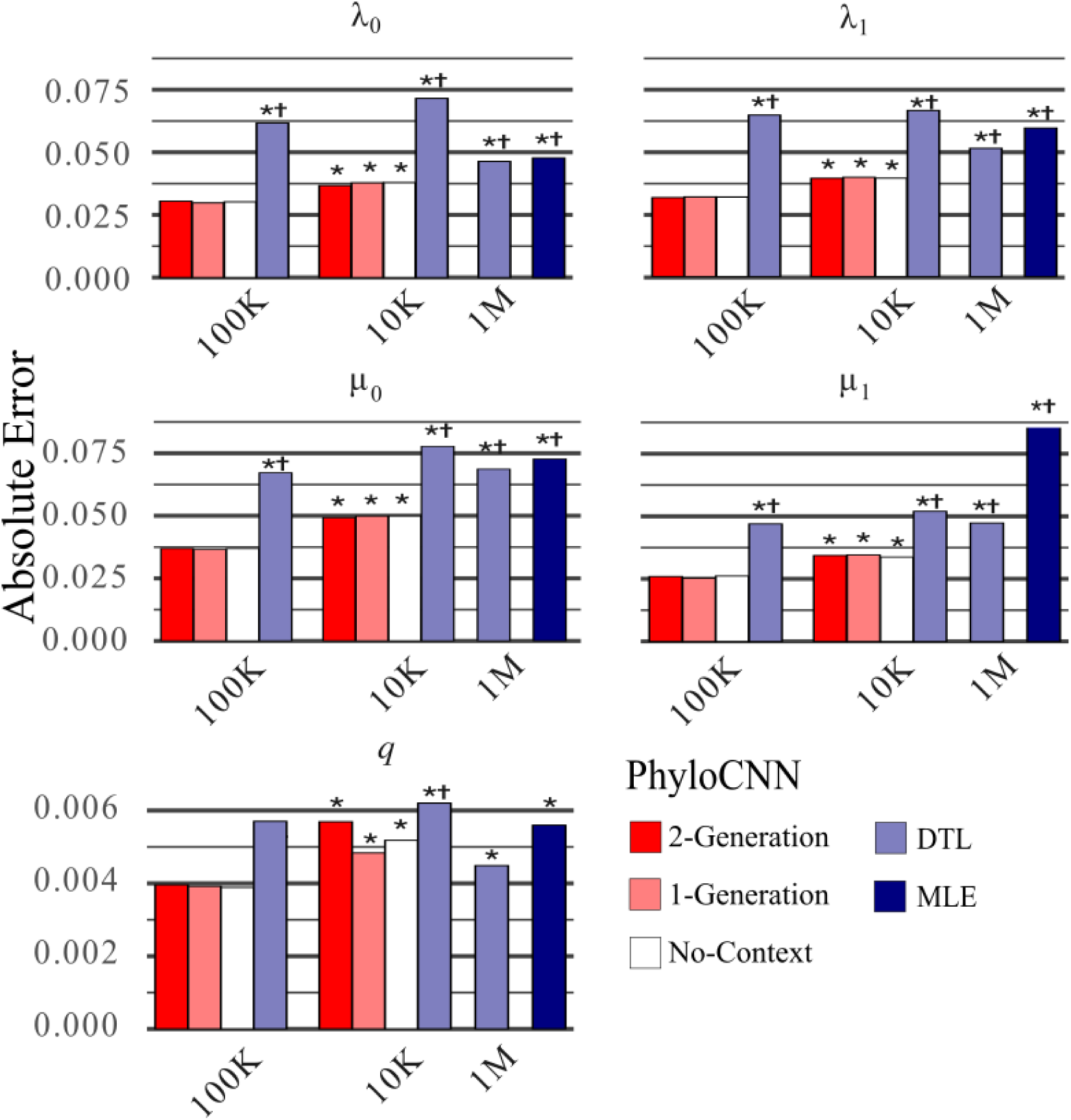
Accuracy of PhyloCNN, DeepTimeLearning (DTL) and diversitree with BiSSE. Accuracy is measured by the mean absolute error (MAE). Results are shown for PhyloCNN with 2-Generation, 1-Generation and No-Context using 100K and 10K training trees, for DTL using CNN and CDV encoding (adapted from CBLV) with 4M, 100K and 10K training trees, and for maximum-likelihood estimation (MLE) implemented in diversitree. Throughout, we used the same 1,000 independent test trees for all methods. Results are shown for parameters estimated under the BiSSE model, with rates associated with character states 0 and 1 for speciation (λ_0_, λ_1_) and extinction (μ_0_, μ_1_), as well as to state transitions occurring anagenetically with symmetric rates (*q*_01_=*q*_10_). **\*** Significantly higher MAE compared to PhyloCNN with 2-Generation and 100K training trees (paired Z-test, p<0.01). † Significantly higher MAE compared to PhyloCNN with 2-Generation and 10K training trees.

We consistently find that the accuracy is higher for each parameter and DL method when the number of training trees is higher (Fig. 5). Obviously, there is no such effect for MLE, which follows a radically different principle without any training phase. We also note that there is no bias tendency for any method (Supp. Tab. 6).

There is almost no difference between the three different generational contexts of PhyloCNN. The small differences observed with 100K training trees are not significant (p>0.01). The only significant difference (p<0.01) with 10K training trees is a higher 2-Generation error for the parameter *q* (= *q*_01_ = *q*_10_), but for this parameter the magnitude of the errors is very low (<1%) and one order of magnitude lower than for the other parameters.

The comparisons between PhyloCNN and DTL confirm the results obtained with phylodynamics models. PhyloCNN with 2-Generation and 100K training trees is significantly (p<0.01) better than DTL trained with 1M trees for all parameters, and the error reduction is ∼50% or more, except for *q* (∼10%). When trained with 10K trees, 2-Generation PhyloCNN is still significantly better than DTL trained with 1M trees for 4 out of 5 parameters (but *q*). The same is true for MLE, which lags behind DTL (Lambert et al. 2023) and thus PhyloCNN for all parameters (Fig. 5). One possible explanation is that MLE is not informed by the priors, as PhyloCNN and DTL are for some of the parameters with non-uniform priors (e.g., λ_1_ and *q*, see above and Supp. Tab. S2).

To summarize, in these results and those obtained with phylodynamics datasets, PhyloCNN requires two orders of magnitude fewer training trees than PhyloDeep using the CBLV (or an adaptation) tree representation (Figs. 4 and 5), and one order of magnitude fewer than when a high-performing set of summary statistics is available and used, as is the case for phylodynamics models but not for BiSSE. This performance of PhyloCNN is the result of (i) a relatively small number of neural network weights (less than FFNN_SumStats, as used in Voznica et al., 2022), and (ii) a good fit between the tree representation and our 2-channel CNN neural architecture. This radical change in computational efficiency and accuracy opens the door to a wide range of applications on new data and models.

### Analysis of the HIV phylodynamics dataset

We analyzed the MSM in Zurich dataset using the dated phylogenetic tree from (Rasmussen et al., 2017), which was also analyzed in (Voznica et al., 2022). We first performed model selection using PhyloCNN with 2-Generation tree representation. PhyloCNN was trained with 10K trees for each of the three models (BD, BDEI, BDSS). These trees were simulated with the same tree size as the HIV tree and the new priors better suited for this dataset (see above; Supp. Tab. S1). In agreement with Rasmussen et al. (2017) and Voznica et al. (2022), the super-spreader BDSS model was selected with an estimated probability of ∼1.00. The *a priori* model adequacy check under BDSS (Supp. Fig. S2) shows that the HIV tree is well included in the PCA cloud of simulated trees, especially when using the new priors and removing the summary statistics related to the sampling over time, which is not completely random due to partner notification (Zhukova and Gascuel, 2024).

We trained PhyloCNN to perform parameter estimation under BDSS with 50K trees that were simulated with the same tree size and the new priors. The computing time to generate and encode these 50K trees and train PhyloCNN on our laptop (Apple M3 Pro with 11 core CPU, 14 core GPU and 16 core Neural Engine) was ∼11 hours, which is still much faster than the time required by BEAST2 for the same dataset (∼80 hours, cf. Voznica et al., 2022). To compare different methods using the new priors, we also trained a FFNN_SumStats network using the same 50K new trees and the same SumStats and neural architecture as for the simulation study (see above). For these two methods, we computed the posterior distribution of the parameters using a form of parametric bootstrap similar to the approach used in PhyloDeep to compute confidence intervals (see Supp. Mat. for details). The results are shown in Figure 6. We also present the results of PhyloDeep with CBLV and BEAST2 reported in (Voznica et al., 2022).

**Figure 6.**
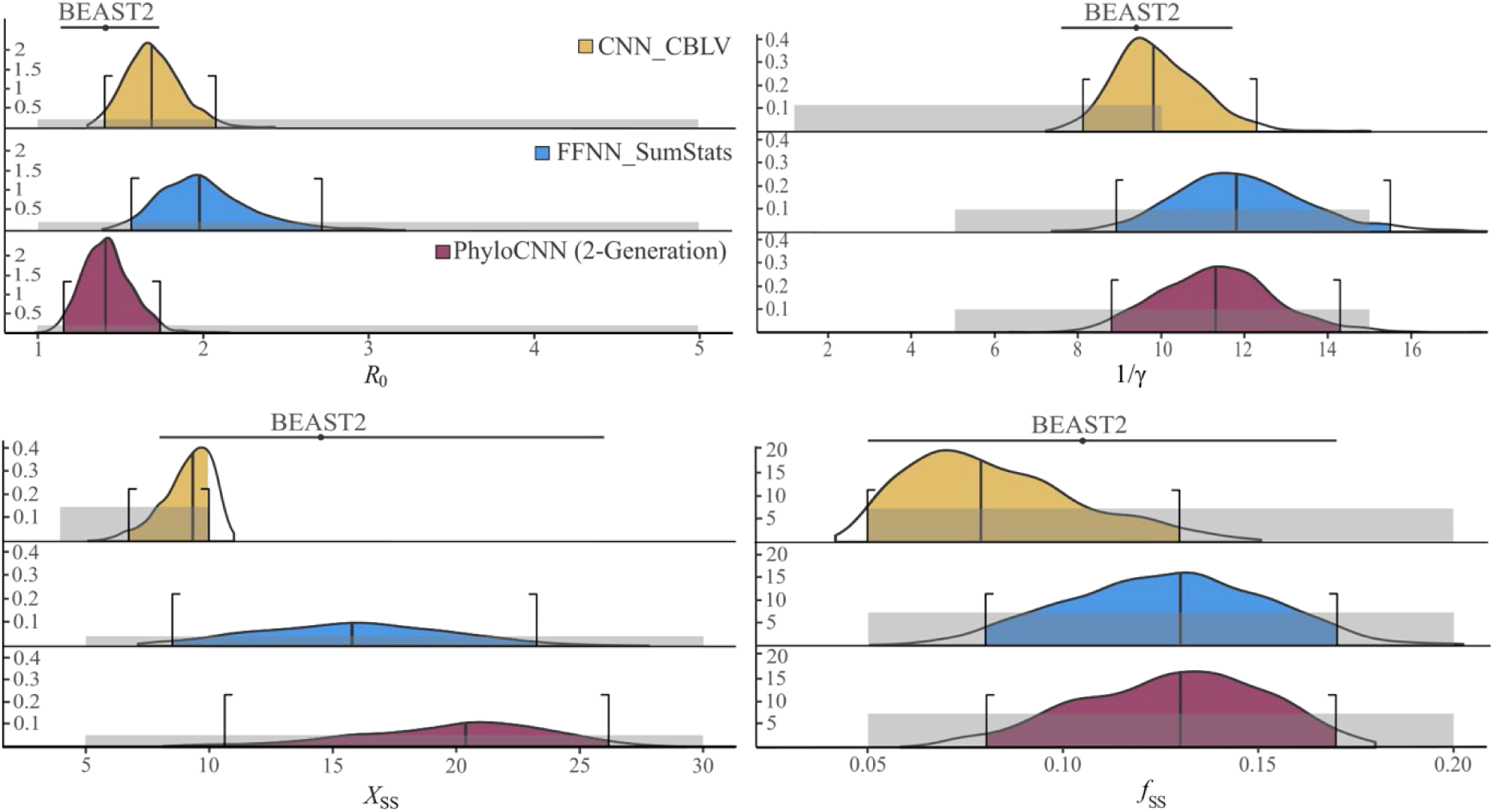
Parameter estimation for the HIV dataset under the BDSS model. Posterior distributions and 95% CIs (colored, within brackets) are shown for *R*_0_, the infectious period 1/γ, the ratio of transmission rates between super- and normal-spreaders *X*_SS_, and the super-spreader fraction *f*_SS_. Network predictions were obtained with CNN_CBLV (yellow, published in Voznica et al., 2022), PhyloCNN trained with specific priors and new simulated trees (2-Generation, purple), and FFNN_SumStats retrained with the new priors and trees (blue). The prior distributions for each parameter and method are shown as gray boxes. We also show the results and 95% CIs for BEAST2 (reported by Voznica et al., 2022).

For all parameters except *R*_0_, the posterior distributions of PhyloCNN and FFNN_SumStats are similar, consistent with the fact that they were trained with the same priors and simulated trees. For *R*_0_, PhyloCNN is more in line with PhyloDeep and BEAST2. In fact, the results of PhyloCNN and BEAST2 are similar for all parameters. In particular, both approaches (and FFNN_SumStats) clearly show the difficulty of estimating the BDSS parameters (*X*_SS_ and *f*_SS_; see also Xie et al., 2024). The confidence intervals with BEAST2 are very large, and the posteriors with PhyloCNN are rather flat and not very informative compared to the priors. PhyloCNN and PhyloDeep predictions are mostly in agreement for the non-BDSS parameters (*R*_0_ and 1/γ), although the prior for 1/γ was different, justifying the new prior used here for this parameter. The main differences between PhyloCNN and PhyloDeep are for the BDSS parameters, due to different priors for *X*_SS_, but also for *f*_SS_ (with identical priors), probably due to the correlation between these two parameters (and the difficulty of estimation). The *a posteriori* model adequacy check consistently shows that the HIV tree is well included in the PCA cloud of the 1K trees we simulated using the posterior distribution of parameters estimated by PhyloCNN, especially when the summary statistics related to sampling over time are removed (Supp. Fig. S3; see Supp. Mat. for details).

Overall, these results demonstrate the usefulness of PhyloCNN in producing consistent results in a relatively low computing time, even when new simulations and training are required due to changes in settings and priors, which is likely to be the norm for most studies and analyses.

### Analysis of the primates diversification dataset

We analyzed the primate dataset using the dated phylogenetic tree of Fabre et al. (2009), which was also analyzed in (Gómez and Verdú, 2012) and (Lambert et al., 2023). We used PhyloCNN with 2-Generation and 100K training trees generated under the BiSSE model, as obtained in the simulation study. PhyloCNN was compared with DeepTimeLearning (DTL) trained with 1M trees and the maximum-likelihood estimation (MLE) approach implemented in diversitree (Fitzjohn, 2012). These two methods were run with the same options and settings as in the simulation study. We did not perform an *a priori* model check, as this was already done in (Lambert et al., 2023) using summary statistics adapted from (Saulnier et al., 2017) and four statistics describing the state distribution (e.g., number of leaves with state 0 vs. 1). An *a posteriori* model check using the same SumStats was performed on the parameter estimates obtained by PhyloCNN, using the same approach as for the HIV tree (Sup. Fig. S4; see Supp. Mat. for details). Results are shown in Figure 7.

**Figure 7.**
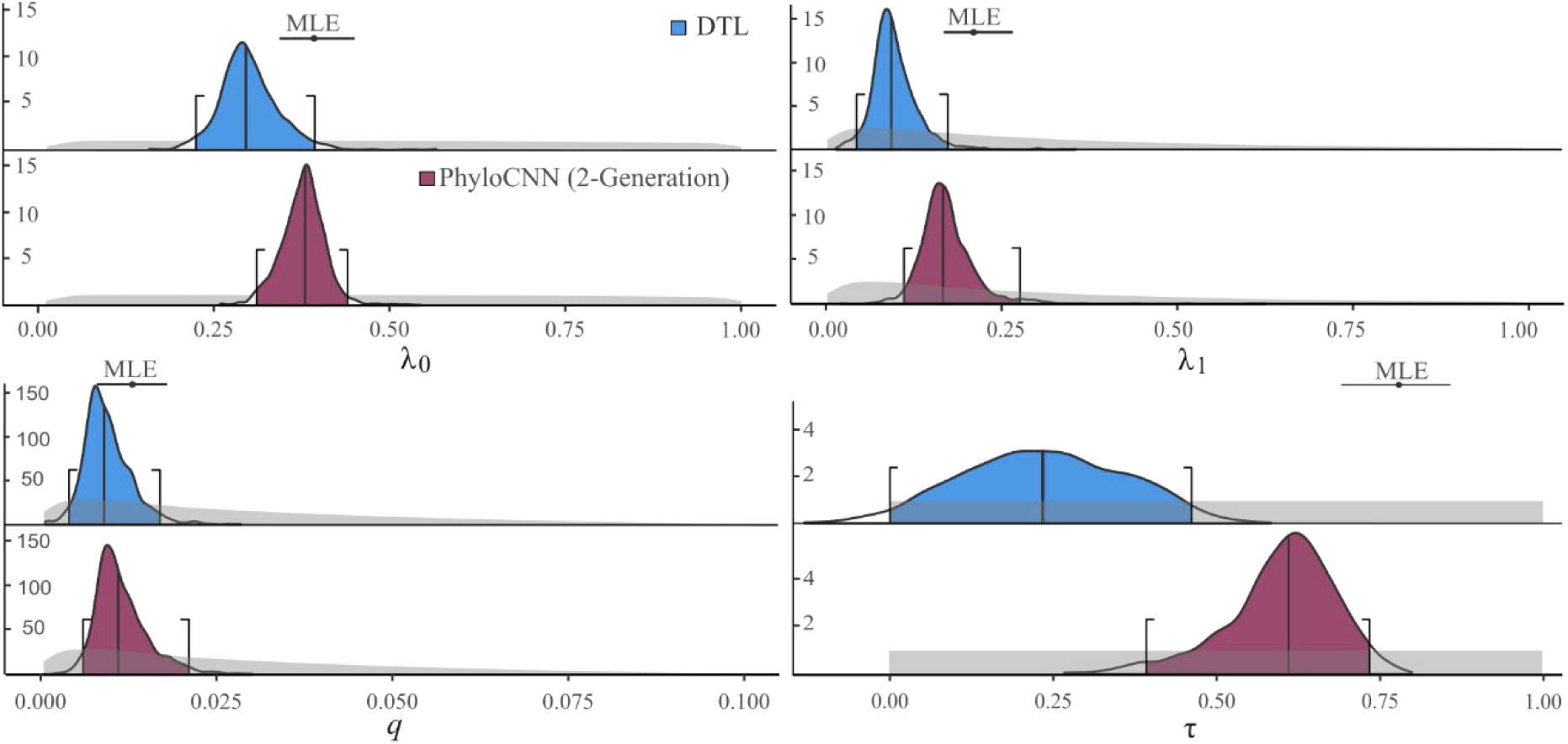
Parameter estimation for the primates dataset under the BiSSE model. Posterior distributions and the 95% CIs (colored, within brackets) are shown for the speciation rates λ_0_ and λ_1_ (mutualistic primates have state 0), for state transition rate *q* (=*q*_01_=*q*_10_) and for turnover τ. We present the results of PhyloCNN (2-Generation, purple) and DeepTimeLearning (blue) using CNN and CDV encoding (adapted from CBLV). The MLE estimate (black dot) and its 95% CI (black bar) are shown above the distributions for each parameter. The prior distributions for each parameter (computed from the 100K training trees used in the simulation study) are shown in gray.

As in (Gómez and Verdú, 2012) and (Lambert et al., 2023), we recover with PhyloCNN that the mutualistic primates (with state 0) have a higher speciation rate than the antagonistic primates (λ_0_ = 0.378 vs. λ_1_ = 0.116). However, for all parameters, the results (point estimates, posterior distributions) with PhyloCNN are significantly closer to the results of MLE than those of DTL. Consistently, the log-likelihood value (computed using diversitree) of DTL parameter estimates is lower than that of PhyloCNN, which is relatively close to the MLE value (−861.7, −850.1 and −846.9, respectively; the difference between MLE and DTL has a p-value of 11.5% using a four-degrees-of-freedom likelihood ratio test, while the difference between MLE and PhyloCNN is negligible). In particular, the PhyloCNN estimate of the turnover parameter τ (= 0.610 [0.39, 0.73]) is consistent with the usual estimates of this parameter for mammals (Stadler and Bokma, 2013), while the DTL turnover estimate (= 0.234 [0.0, 0.46]) seems too low. The *a posteriori* adequacy model check also indicates the consistency of the PhyloCNN estimates (Supp. Fig. S4). Overall, the results of PhyloCNN are in agreement with previous studies and estimates and are quite credible when we refer to its accuracy with simulations.

## Discussion

Our tree representation based on node features and neighborhoods offers a drastic reduction in the number of simulated trees needed to train convolutional network architectures compared to our previous CBLV representation (Voznica et al., 2022) and its adaptations (Lambert et al., 2023; Thompson et al., 2024). CBLV requires millions of simulated trees, which is memory and computationally expensive, whereas PhyloCNN achieves similar or better accuracy with a few tens of thousands of trees (Figs. 4, 5, Sup. Tab. S3). When an extensive set of informative summary statistics is available, as in phylodynamics (Saulnier et al., 2027; Voznica et al., 2022), the difference between PhyloCNN and a simple FFNN_SumStats architecture is still noticeable, but less pronounced. However, for some models, in particular BiSSE (Maddison et al., 2007) and MuSSE (Fitzjohn, 2012), which are commonly used in macroevolution and phylogeography, we do not have such summary statistics (Janzen and Etienne, 2024), and having a simple, complete, and general tree representation like PhyloCNN’s is a major advantage. This advantage is clearly illustrated by the application of PhyloCNN under BiSSE to the primate dataset, where our results are close to maximum-likelihood estimates and similar to general estimates obtained for mammals (Stadler and Bokma, 2013). This achievement, as well as the low computational cost of the HIV study with specific simulations and priors, is encouraging to ensure that PhyloCNN can be successfully applied to the complex datasets collected in phylodynamics and diversification studies.

The gain in accuracy of PhyloCNN is especially sensible with the most complex models, in particular with BiSSE (Fig. 5). The factors are certainly multiple, but part of the explanation for the BiSSE model (and by extension MuSSE) probably comes from the processing of trait states in ancestral tree nodes. In the PhyloCNN tree representation, the number of node descendants in state 0 vs. 1 is explicitly indicated. This somehow allows for ancestral reconstruction of node states with high accuracy (Gascuel and Steel, 2014), whereas this information is very difficult to extract in the CBLV representation. A more general explanation relates to the simultaneous design of the tree representation and the neural architecture. To develop PhyloCNN, we experimented with different approaches, such as graph neural networks (GNNs) with differentiable pooling (Ying et al., 2018), as in the approach applied to ancestral recombination graphs in population genetics by Korfmann et al. (2024). This resulted in poorer accuracy than CBLV, likely caused by oversmoothing due to the sparsity of the “graph” (i.e., tree), consistent with the observations of Lajaaiti et al. (2023). Our tree representation is specifically designed for a CNN architecture and follows the same rationale as the sliding windows used in image analysis. A key feature is the two different channels for internal nodes and tree leaves. The measurements associated with each node (e.g., under BiSSE, the number of descendants in state 0 vs. 1) have also been explored and tuned, although some progress could probably still be made. Importantly, the PhyloCNN architecture involves a relatively small number of neural network weights (less than PhyloDeep with summary statistics), which likely explains its accuracy with a small number of training trees. All these features should guide future methodological developments and facilitate application to a wide variety of datasets, models, and questions in evolutionary biology.

Several avenues for further research are open: (1) Analysis of large trees, e.g. using subtrees, as performed by PhyloDeep with phylodynamics models (Voznica et al., 2022). (2) Estimating parameter changes over time and along the input phylogenetic tree, as in the Bayesian skyline plot approach (Stadler et al., 2012). (3) For multistate and multitype models (e.g., BiSSE, MuSSE, BDSS, and MTBD, Stadler and Bonhoeffer, 2013), estimating the posterior of the state/type for each tree node and leaf (e.g., super-vs. normal-spreader). (4) Improving the robustness of network predictions in the presence of model and simulation misspecifications and approximations, e.g., using more realistic simulated training trees (Xie et al., 2024) or domain adaptation techniques from machine learning (Mo and Siepel, 2023). (5) Finally, all these developments should be incorporated into a library or pipeline, such as phyddle (Landis and Thompson, 2024), to help inexperienced users build their own deep learning tools and study their favorite datasets, phylogenetic trees and models.

## Availability

Instructions to install PhyloCNN, as well as the trees used as test sets, all scripts and notebooks are available from https://github.com/manolofperez/phyloCNN. HIV and primates datasets are available from https://doi.org/10.5061/dryad.prr4xgxx9. See Supplementary Material for details.

## Acknowledgements

We thank Anna Zhukova, Jakub Voznica, Ruopeng Xie and Frédéric Lemoine for their help with tree simulators, implementation and comments on the manuscript, and Flora Jay and Vinh-Son Pho for their suggestions on deep learning.

## SUPPLEMENTARY MATERIAL

### Material and Methods

#### Simulations

- We performed phylodynamics simulations using the simulators provided by Voznica et al. (2022), available at: https://github.com/evolbioinfo/phylodeep/tree/main/simulators/bd_models. We wrote a Python script to draw parameter samples from the prior distribution, and to format the sampled parameters for input to the simulators. The following commands were used in Python 3.9.18 with the PhyloCNN conda environment (available on GitHub):

- Sample from prior distributions by specifying the model, the minimum and maximum bounds of each parameter, the number of simulations, and the name of the output file. BD model (-m=model; -r=*R*_0_; -i=1/γ; -s=tree size; -p=sampling probability; -n=number of samples; -o: output file): `python generate_parameters.py -m BD -r 1,5 -i 1,10 -s 200,500 -p 0.01,1 -n 10000 -o parameters_BD.txt` BDEI model (-m=model; -r=*R*_0_; -i=1/γ; -e=incubation factor (ε/γ); -s=tree size; -n=number of samples; -p=sampling probability; -o: output file): `python generate_parameters.py -m BDEI -r 1,5 -i 1,10 -e 0.2,5 -s 200,500 -p 0.01,1 -n 10000 -o parameters_BDEI.txt` BDSS model (-m=model; -r=*R*_0_; -i=1/γ; -x=*X*_SS_; -f=*f*_SS_; -s=tree size; -p=sampling probability; -n=number of samples; -o: output file): `python generate_parameters.py -m BDSS -r 1,5 -i 1,10 -x 3,10 -f 0.05,0.2 -s 200,500 -p 0.01,1 -n 10000 -oparameters_BDSS.txt`
- Simulate trees using the Python scripts from Voznica et al. (2022). The example below is for BD, but the same command was used for other models. It requires the simulator to be called along with the parameter file generated in the previous step (e.g., parameters_BD.txt) and the maximum simulation time (with a default of 500; Voznica et al., 2022): `python TreeGen_BD_refactored.py parameters_BD.txt` `<max_time=500> > BD_trees.nwk`
- BiSSE simulations were carried out with the R package castor v1.6.6 (Louca and Doebeli, 2018) using the script provided by Lambert et al. (2023), available at: https://github.com/JakubVoz/deeptimelearning/tree/main/simulators/BiSSE. To run it, we used the Python script generate_parameters.py to draw samples from the prior distribution and format the sampled parameters for input to the simulator (-m=model; -l0 =λ_0_; -t=τ; -l1=ratio between λ_1_ and λ_0_; -q=ratio between *q* (= *q*_01_ = *q*_10_) and λ_0_; -s=tree size; -n=number of samples; -p=sampling probability; -o=output file): `python generate_parameters.py -m BISSE -l0 0.01,1.0 -t 0,1 -l1 0.1,1.0 -q 0.01,0.1 -s 200,500 -p 0.01,1 -n 10000 -oparameters_BiSSE.txt`

- The simulation is then performed using the script from Lambert et al. (2023) in R v4.4.1 (along with parameters required by the script, such as the indice, seed number, step, number of retrials, and output file names). The values used are indicated below: `Rscript BiSSE_simulator.R parameters_BiSSE.txt <indice=1> <seed_base=12345> <step=10> <nb_retrials=100> BiSSE_trees.nwk BiSSE_stats.txt BiSSE_params.txt`

#### Encoding

- Encoding was performed with the script PhyloCNN_Encoding_PhyloDyn.py for all phylodynamics models (BD, BDEI, and BDSS). The script requires the input tree file (-t) and the output file (-o). The output contains the encoding information for each leaf and internal node of each tree (rescaled to an average branch length equal to 1.0, as described in the *Encoding phylogenies based on node neighborhood* section) and the rescaling factor for that tree. `python PhyloCNN_Encoding_PhyloDyn.py -t BD_trees.nwk -o Encoded_trees_BD.csv`
- A similar process was used for BiSSE, using a specific encoding script for this model (PhyloCNN_Encoding_BiSSE.py) that includes information about leaf states and number of leaves in state 0 for internal nodes. This script also requires the input tree file (-t) and the output file (-o). As in the phylodynamics case, the script outputs the encoding information for each leaf and internal node for each tree (rescaled to an average branch length equal to 1.0, as described in the *Encoding phylogenies based on node neighborhood* section) and the rescaling factor for that tree. `python PhyloCNN_Encoding_BiSSE.py -t BiSSE_trees.nwk -o Encoded_trees_BiSSE.csv`

#### Preprocessing encoded trees, Training PhyloCNN, and Predicting the test sets

- We performed (1) preprocessing, (2) training, and (3) predictions of the test sets using jupyter notebooks that are annotated and show examples of the expected outputs to ensure reproducibility.

- For preprocessing, we import the parameter file generated in the simulations (e.g., parameters_BD.txt) and scale the time-dependent parameters by dividing (1/γ and 1/ε for phylodynamics models; Voznica et al., 2022) or multiplying them (λ_0_, λ_1_, μ_0_ and μ_1_ for BiSSE; Lambert et al., 2023) according to the rescaling factor used for encoding the trees. Then, we import the encoded trees (e.g., Encoded_trees_BD.csv) and add the sampling probability as an additional feature to the encoding. We order the nodes according to their distance to the root and split leaves and internal nodes in separate tables (channels) to be input in PhyloCNN for training.
- We trained PhyloCNN to perform both model selection (in phylodynamics, using input trees from all three models: BD, BDEI, and BDSS) and parameter estimation under each birth-death model (BD, BDEI, BDSS, and BiSSE).
- The trained PhyloCNN networks were then used to perform predictions on the test sets to evaluate accuracy and compare the performance with other methods.
- Specifically, the Github repository contains notebooks for preprocessing, training, and performing predictions for:

- Model selection - PhyloCNN_Train_PhyDyn_ModelSelection.ipynb;
- Parameter estimation for the BD model - PhyloCNN_Train_BD.ipynb;
- Parameter estimation for the BDEI model - PhyloCNN_Train_BDEI.ipynb;
- Parameter estimation for the BDSS model - PhyloCNN_Train_BDSS.ipynb;
- Parameter estimation for the BiSSE model - PhyloCNN_Train_BiSSE.ipynb.

#### Confidence intervals (CI) and posterior distributions with empirical datasets

As in (Voznica et al., 2022), we used an approximate version of the parametric bootstrap to compute the posterior distributions and 95% confidence intervals (CIs) of the parameter estimates. This approach is based on using the distribution of prediction errors from simulations in the training set that are closest to the empirical data and tree *T*, thereby avoiding the need for new simulations as required by the standard parametric bootstrap. To do this, we used the trained network to make predictions on the training simulations (100K trees for the primates dataset with BiSSE, and 50K trees for the HIV dataset with BDSS). To build the posterior distributions and CIs, for each training simulation that is close enough to *T,* we compared the true parameter values (used to simulate the tree) and the corresponding predicted parameter values obtained from the trained network. We provide notebooks to reproduce these computations for the HIV (CI_HIV.ipynb) and primates (CI_primates.ipynb) datasets. This includes the following steps:

- In the first step, the simulations closest to *T* are selected based on tree size and sampling probability. This procedure was not applied to the BDSS simulations for HIV, which were run with customized priors (Supp. Tab. S1), a fixed tree size (= 200 tips), and a reduced sampling probability interval (uniform from 0.2 to 0.3). For BiSSE and the primates, the selection procedure was as follows:

- Tree size: We first select the 20% of simulations (out of 100K, i.e. 20k) that are closest to *T* in terms of tree size (= 260 tips).
- Sampling probability: From this subset, we select the 20% of simulations that are closest to *T* in terms of sampling probability, i.e. 4,000 simulations.
- The second step was used for both HIV and primates analyses. It includes: identifying nearest neighboring parameter values, calculating estimation errors, and centering errors around parameter predictions:

- For each model parameter with value *p* predicted from *T*, we select the 1000 simulations from the (reduced to 4K with primates; 50K with HIV) training set whose true parameter values *r*_i_ are closest to *p*. We record these true parameter values *R* = {*r_i_*_=1,1000_} and their corresponding predicted values *P* = {*p_i_*_=1,1000_}.
- The estimation errors for these neighboring simulations are computed as *E* = {*e_i_*=*p_i_*−*r_i_*}.
- To adjust for bias, we center the errors around *p* by subtracting the median of errors *m*(*E*), resulting in the posterior distribution *D* = {*p*+*e*_i_− *m*(*E*)}. Negative values in *D* are set to zero.
- Extracting CIs and posterior distributions:

- We extract the 95% CI around *p* from the posterior distribution *D* by determining the 2.5th and 97.5th percentiles.
- The obtained CIs and posterior distributions *D* are shown in Figure 6 (for the HIV dataset) and Figure 7 (for the primates dataset).

#### Prior and posterior model adequacy checks

*A priori* model adequacy checks were already performed in Voznica et al. (2022) for the HIV dataset under BDSS, and in Lambert et al. (2023) for the primates dataset under BiSSE. Since we changed the priors for analysing the HIV dataset and simulated new BDSS trees based on these new priors and having the same size (= 200 tips) as the HIV tree, we repeated this procedure. We performed a principal component analysis (PCA) on the SumStats of both simulations and the HIV tree, and we evaluated each of the SumStats by checking whether the corresponding value for the HIV tree was between the minimum and maximum values observed in the simulated trees (Supp. Fig. S2). The implementation is as follows:

- We computed the SumStats of the HIV tree and of 1000 BDSS trees using scripts provided in https://github.com/evolbioinfo/phylodeep/blob/main/phylodeep/sumstats.py. We reduced and centered the SumStats (using sklearn.preprocessing.StandardScaler) and performed a Principal Component Analysis (using sklearn.decomposition.PCA) in scikit-learn v1.5.1. An empirical dataset positively passes the check when it is inside the cloud of simulated data.
- We also evaluated the SumStats individually. For each of the summary statistics, we checked whether the value obtained for the HIV dataset was between the minimum and maximum values of that summary statistic in 10K new simulated trees. For both empirical datasets (HIV and primates), we also performed sanity checks of the posterior distributions obtained with PhyloCNN. We simulated 1000 new trees under BDSS for HIV and BiSSE for the primates. The parameter values for the simulations were sampled from their respective PhyloCNN posterior distributions (shown in Fig. 6 for HIV, and Fig. 7 for primates). The sampling probability was drawn uniformly between 0.2 and 0.3 for HIV and equal to 0.68 for the primates dataset. We then performed the same two complementary steps as in the *a priori* check:
- PCA (SumStats for BiSSE were obtained using the notebook BiSSE_SumStats.ipynb, modified from Lambert et al. 2023).
- Individual evaluation of the SumStats to check that the observed values are between the minimum and maximum values of the 1000 new posterior-based simulations.

**Table S1:** Priors for phylodynamics birth-death models.

| Parameters | Symbol | Parameter Relationships | Range Simul. Study | Range HIV dataset |
| --- | --- | --- | --- | --- |
| Tree size | $t$ | | $U(200, 500)$ | 200 |
| Sampling probability | $s$ | | $U(0.01, 1.0)$ | $U(0.2, 0.3)$ |
| Basic reproductive number | $R_0$ | $=\beta/\gamma$ (BD and BDEI)<br>$= (\beta_{S,S}+\beta_{N,N})/\gamma$ (BDSS) | $U(1.0, 5.0)$ | Same |
| Infectious period | $1/\gamma$ | | $U(1.0, 10.0)$ | $U(5.0, 15.0)$ |
| Incubation factor<sup>1</sup> | $f_i$ | $=\varepsilon/\gamma$ | $U(0.2, 5.0)$ | Same |
| Incubation period<sup>1</sup> | $1/\varepsilon$ | $=1/f_i*\gamma$ | $[0.2, 50.0]$ | Same |
| Superspreading infectious ratio<sup>2</sup> | $X_{SS}$ | $=\beta_{S,S}/\beta_{N,S}=\beta_{S,N}/\beta_{N,N}$ | $U(3.0, 10.0)$ | $U(5.0, 30.0)$ |
| Fraction of superspreading individuals<sup>2</sup> | $f_{SS}$ | $=\beta_{S,S}/(\beta_{S,S}+\beta_{S,N})$<br>$=1-\beta_{N,N}/(\beta_{N,N}+\beta_{N,S})$ | $U(0.05, 0.2)$ | Same |

Parameters specific to <sup>1</sup>BDEI and <sup>2</sup>BDSS models.

BD parameters are common to all three models. BDEI-specific parameters^1^ and BDSS-specific parameters^2^ are indicated by superscripts. For the simulation study, we used the same uniform distributions and ranges as in (Voznica et al., 2022). For the analysis of the HIV dataset, we changed the ranges and priors of some the parameters (see corresponding subsection in the main manuscript). All parameters were sampled from uniform distributions, denoted by *U*(*x*, *y*), where *x* is the lower bound and *y* is the upper bound of the distribution. The incubation period (1/ε) was not sampled, its value depends on *f*_I_ and γ, and its distribution is non-uniform with the range indicated between brackets. See Fig. 1 for more information on these models, and (Voznica et al., 2022) for explanations on model parametrization.

**Table S2:** Priors for the BiSSE model.

| Parameters | Symbol | Parameter Relationships | Range |
| --- | --- | --- | --- |
| Tree size | $t$ | | $U(200, 500)$ |
| Sampling probability | $s$ | | $U(0.01, 1.0)$ |
| Turnover rate | $\tau$ | | $U(0.0, 1.0)$ |
| Speciation rate state 0 | $\lambda_0$ | | $U(0.01, 1.0)$ |
| Ratio of speciation rates of states 1 and 0 | $r_\lambda$ | | $U(0.1, 1.0)$ |
| Ratio of transition rate to speciation rate of state 0 | $r_q$ | | $U(0.01, 0.1)$ |
| Speciation rate of state 1 | $\lambda_1$ | $= \lambda_0 * r_\lambda$ | $[10e-3, 1.0]$ |
| Transition rate | $q = q_{01} = q_{10}$ | $= \lambda_0 * r_q$ | $[10e-4, 0.1]$ |
| Extinction rate state 0 | $\mu_0$ | $= \tau * \lambda_0$ | $[0.0, 1.0]$ |
| Extinction rate state 1 | $\mu_1$ | $= \tau * \lambda_1$ | $[0.0, 1.0]$ |

We used the same priors as (Lambert et al., 2023). For each parameter, we present the range of values used in the simulations for the training, validation, and testing sets. We used uniform distributions, denoted by *U*(*x*, *y*), where *x* and *y* are the lower and upper bounds, respectively. *λ*_1_ and *q* (= *q*_01_ = *q*_10_) are parameterized relative to *λ*_0_. The extinction rates (μ_0_ and μ_1_) are not estimated by the network, but are calculated using by their respective speciation rate (*λ*_0_ and *λ*_1_) and the turnover (τ). The range of values for parameters calculated relative to others is given in brackets. If in a given simulation, the tree leaves with state 1 are more frequent than the leaves with state 0, the state values are inverted and the parameter values are changed accordingly; for example, λ_1_ consistently becomes λ_0_ and vice versa. This explains why we can observe (∼20% of the samples in our experiments) λ_1_ > λ_0_.

**Table S3:** Running times and storage required for DL methods.

| Step | Time in hours | Storage |
| --- | --- | --- |
| Simulating trees, phylodynamics<br>BD, BDEI, and BDSS models<sup>1</sup> | ~13h for 100K trees | ~500 MB for 100K trees |
| Simulating BiSSE trees<sup>1</sup> | ~4h for 100K trees | ~700 MB for 100K trees |
| PhyloCNN tree encoding<sup>1</sup><br>2-Generation | ~2h for 100K trees | ~10GB for 100K trees |
| CBLV tree encoding<sup>1</sup> | ~1.5h for 100K trees | ~1GB for 100K trees |
| Training PhyloCNN (50K trees)<sup>1</sup><br>HIV dataset | ~3.5h | - |
| Training PhyloCNN (100K trees)<sup>2</sup><br>Simulation study | ~2h | - |
| Training PhyloDeep<br>FFNN_SumStats (4M trees)<sup>3</sup> | ~5h | - |
| Training PhyloDeep<br>CNN_CBLV (4M trees, BDSS)<sup>3</sup> | ~150h | - |
| Training DTL (1M trees)<sup>3</sup> | ~8h | - |

<sup>1</sup>Apple M3 Pro (11 core CPU, 14 core GPU, 16 core Neural Engine); <sup>2</sup>Nvidia Tesla V100 GPUs;
<sup>3</sup>Nvidia Titan X GPUs (Voznica et al., 2022; Lambert et al., 2023).

The simulation scripts for PhyloCNN are the same as for PhyloDeep (phylodynamics) and DTL (BiSSE). PhyloCNN encoding takes slightly more time per tree. This is compensated by the much smaller size (40 times smaller than for PhyloDeep and 10 times smaller than for DTL) of the training sets required to achieve similar or better accuracy than CNN_CBLV (Supp. Tabs. S4-S6).

We provide the storage required for simulated and encoded trees. Again, PhyloCNN uses at least ten times fewer trees for training, resulting in a much smaller storage requirement. PhyloCNN encoding has a much higher storage requirement per tree (especially when using the 2-Generation context). However, this difference is compensated by the smaller number of training trees for PhyloCNN. In addition, the memory requirement of PhyloCNN can be bypassed because trees can be encoded on the fly during the learning phase.

We provide the runtimes for training PhyloCNN with 100K trees (as used in Figs. 4-5 and Supp. Tabs. S4 and S6; Nvidia Tesla V100 GPUs). Compared to PhyloDeep (4M training trees) and DTL (1M training trees), PhyloCNN was always faster. Although we used GPUs of different generations (the Nvidia Tesla V100 used for PhyloCNN is ∼60% faster than the Nvidia Titan X used for the other DL approaches), the difference in running times is still substantial (to very high for BDSS with CNN_CBLV).

To predict the HIV dataset, we trained PhyloCNN with 50K trees simulated under different priors (Supp. Tab. S1). To demonstrate the potential of PhyloCNN to adapt to new datasets, models, and priors, even in an accessible computational environment, all computations were performed on a MacBook Pro laptop instead of using a powerful computer cluster. The total running time was ∼11h to analyze this dataset, compared to 80h for BEAST2 (see text for details and references).

**Table S4:** Phylodynamics models, results with 100K training trees (and more)

|  |  | Mean Relative Error (Mean Relative Bias) |  |  |  |  |  |  |
| --- | --- | --- | --- | --- | --- | --- | --- | --- |
| Model |  | PhyloCNN<br>(100K training trees) |  |  | PhyloDeep<br>(100K training trees) |  | PhyloDeep<br>(4M training trees) |  |
| BD | Parameter | 2-Generation | 1-Generation | No-Context | FFNN_SumStat | CNN_CBLV | FFNN_SumStat | CNN_CBLV |
| | $R_0$ | 0.070 (-0.004) | 0.069 (-0.007) | 0.069 (-0.009) | 0.074 (-0.019) | 0.082* (-0.004) | 0.070 (-0.012) | 0.075* (-0.010) |
| | $1/\gamma$ | 0.065 (-0.010) | 0.064 (-0.009) | 0.064 (-0.011) | 0.067 (-0.018) | 0.080* (-0.014) | 0.065 (-0.011) | 0.066 (-0.007) |
| BDEI | $R_0$ | 0.078 (-0.016) | 0.076 (-0.008) | 0.080 (0.025) | 0.080 (-0.007) | 0.100* (-0.024) | 0.079 (-0.018) | 0.082 (-0.021) |
| | $1/\gamma$ | 0.085 (-0.001) | 0.085 (-0.032) | 0.089 (-0.024) | 0.095* (0.000) | 0.129* (-0.018) | 0.092* (-0.017) | 0.092* (-0.005) |
| | $1/\varepsilon$ | 0.105 (-0.009) | 0.110 (-0.016) | 0.130* (0.013) | 0.118* (-0.004) | 0.163* (-0.021) | 0.115* (-0.017) | 0.123* (-0.010) |
| BDSS | $R_0$ | 0.084 (-0.008) | 0.084 (-0.007) | 0.084 (-0.008) | 0.088 (-0.002) | 0.102* (-0.012) | 0.085 (-0.006) | 0.089 (-0.007) |
| | $1/\gamma$ | 0.077 (-0.002) | 0.077 (-0.004) | 0.078 (-0.002) | 0.081 (-0.007) | 0.093* (-0.018) | 0.078 (-0.007) | 0.078 (-0.006) |
| | $X_{ss}$ | 0.128 (-0.003) | 0.130 (-0.006) | 0.135 (-0.038) | 0.139* (-0.013) | 0.170* (-0.008) | 0.135 (-0.010) | 0.139* (0.002) |
| | $f_{ss}$ | 0.239 (-0.022) | 0.240 (-0.023) | 0.252 (0.047) | 0.260* (-0.023) | 0.296* (-0.036) | 0.255* (-0.067) | 0.258* (-0.022) |

\*Significantly higher relative error compared to 2-Generation PhyloCNN (paired Z-test; $p < 0.01$ ).

MRE and MRB values are shown for PhyloCNN with 2-Generation, 1-Generation and No-Context, using 100K training trees. We compared these results with those of the two versions of PhyloDeep, CNN_CBLV and FFNN_SumStats, using 4M and 100K training trees. We used the same 1000 test trees for all methods, taken from (Voznica et al., 2022). The MRE results shown here are also shown in Figure 4.

Among the three options of PhyloCNN, No-Context shows the worst accuracy compared to 1-Generation and 2-Generation, although the differences are not significant (except for 1/ε with BDEI).

When we compare PhyloCNN (2-Generation) with the other DL approaches, it consistently outperforms CNN_CBLV trained with the same number of trees (100K), with significant differences in accuracy for all models and parameters (p<0.01). This better accuracy also holds when PhyloCNN (2-Generation) is compared to CNN_CBLV trained with 4M trees (all differences are positive, 5 out of 9 are significant, p<0.01).

PhyloCNN with 2-Generation also outperforms FFNN_SumStats when trained with the same number of trees (100K) for all models and parameters, but to a lesser extent (all differences are positive, 4 out of 9 are significant, p<0.01). Compared to FFNN_SumStats trained with 4M trees, PhyloCNN with 2-Generation and 100K training trees is better or equal for all models and parameters (all differences are positive or null, 3 out of 9 are significant, p<0.01).

These results also show that the bias (in parentheses) induced by these different methods is small compared to the prediction errors.

**Table S5:** Phylodynamics models, results with 10K training trees.

|  |  | Mean Relative Error (Mean Relative Bias) |  |  |  |  |
| --- | --- | --- | --- | --- | --- | --- |
| Model |  | PhyloCNN<br>(10K training trees) |  |  | PhyloDeep<br>(10K training trees) |  |
| BD | Parameter | 2-Generation | 1-Generation | No-Context | FFNN_SumStats | CNN_CBLV |
| | $R_0$ | 0.076 (-0.028) | 0.075 (-0.029) | 0.075 (-0.027) | 0.080 (-0.006) | 0.105* (-0.045) |
| | $1/\gamma$ | 0.072 (-0.023) | 0.070 (-0.027) | 0.065 (0.002) | 0.078 (-0.007) | 0.112* (-0.040) |
| BDEI | $R_0$ | 0.083 (-0.014) | 0.084 (-0.025) | 0.087 (-0.024) | 0.086 (-0.009) | 0.115* (0.026) |
| | $1/\gamma$ | 0.097 (-0.014) | 0.102 (-0.023) | 0.113* (0.022) | 0.109* (-0.008) | 0.191* (0.106) |
| | $1/\epsilon$ | 0.127 (-0.016) | 0.133 (-0.021) | 0.153* (-0.088) | 0.132 (0.009) | 0.207* (-0.023) |
| BDSS | $R_0$ | 0.090 (-0.004) | 0.091 (-0.015) | 0.088 (-0.003) | 0.100* (-0.005) | 0.130* (-0.065) |
| | $1/\gamma$ | 0.085 (-0.002) | 0.086 (-0.015) | 0.084 (0.006) | 0.094* (-0.006) | 0.129* (-0.037) |
| | $X_{ss}$ | 0.144 (-0.009) | 0.155* (-0.019) | 0.156* (-0.008) | 0.154* (-0.011) | 0.205* (-0.005) |
| | $f_{ss}$ | 0.251 (-0.050) | 0.262 (-0.017) | 0.284* (0.013) | 0.286* (-0.043) | 0.338* (-0.051) |

\*Significantly higher relative error compared to 2-Generation PhyloCNN (paired Z-test; $p < 0.01$ ).

We compared the results of PhyloCNN with those of the two PhyloDeep versions, CNN_CBLV and FFNN_SumStats, also trained on the same 10K trees. We used the same 1000 test trees for all methods, taken from (Voznica et al., 2022). The MRE results shown here are also shown in Figure 4.

Among the three options of PhyloCNN, No-Context shows the worst accuracy, especially for the most complex models (BDEI, BDSS) and the most difficult to predict parameters (1/ε, *X*_SS_, *f*_SS_). The differences between 1-Generation and 2-Generation are small, with 1-Generation being significantly worse only for *X*_SS_ (p<0.01).

When we compare PhyloCNN with 2-Generation to the other DL approaches, it significantly outperforms CNN_CBLV for all models and parameters (p<0.01). PhyloCNN with 2-Generation also outperforms FFNN_SumStats in most cases (differences in accuracy are significant for 5 out of 9 comparisons, p<0.01).

These results also show that the bias (in parentheses) induced by these different methods is small compared to the prediction errors.

**Table S6:** BiSSE diversification model, comparison of methods.

|  | Mean Absolute Error (Mean Bias) |  |  |  |  |  |  |  |  |  |
| --- | --- | --- | --- | --- | --- | --- | --- | --- | --- | --- |
|  | PhyloCNN (100K trees) |  |  | PhyloCNN (10K trees) |  |  | DTL |  |  | MLE |
| Parameter | 2-Generation | 1-Generation | No-Context | 2-Generation | 1-Generation | No-Context | 1M trees | 100K trees | 10K trees |  |
| $\lambda_0$ | 0.031<br>(-0.004) | 0.030<br>(0.005) | 0.030<br>(-0.002) | 0.037*<br>(-0.002) | 0.038*<br>(0.001) | 0.038*<br>(-0.012) | 0.046*†<br>(0.005) | 0.062*†<br>(0.011) | 0.072*†<br>(-0.039) | 0.048*†<br>(0.022) |
| $\lambda_1$ | 0.031<br>(-0.004) | 0.032<br>(0.003) | 0.031<br>(-0.007) | 0.039*<br>(-0.012) | 0.039*<br>(-0.002) | 0.039*<br>(-0.008) | 0.051*†<br>(-0.012) | 0.064*†<br>(-0.034) | 0.066*†<br>(-0.031) | 0.059*†<br>(0.025) |
| $q$ | 0.0040<br>(0.0003) | 0.0039<br>(0.0002) | 0.0039<br>(-0.0003) | 0.0057*<br>(0.0027) | 0.0048*<br>(0.0009) | 0.0052*<br>(-0.0025) | 0.0045*<br>(0.0009) | 0.0057*<br>(0.0004) | 0.0062*<br>(0.0007) | 0.0056*<br>(0.0028) |
| $\mu_0$ | 0.036<br>(0.000) | 0.036<br>(0.004) | 0.036<br>(-0.005) | 0.049*<br>(-0.001) | 0.049*<br>(-0.001) | 0.049*<br>(-0.007) | 0.068*†<br>(0.028) | 0.066*†<br>(0.013) | 0.077*†<br>(-0.039) | 0.072*†<br>(0.050) |
| $\mu_1$ | 0.025<br>(-0.001) | 0.025<br>(0.002) | 0.026<br>(-0.007) | 0.034*<br>(-0.005) | 0.034*<br>(-0.002) | 0.033*<br>(-0.005) | 0.046*†<br>(-0.012) | 0.046*†<br>(-0.016) | 0.051*†<br>(-0.027) | 0.088*†<br>(0.064) |

\*Significantly higher absolute error compared to 2-Generation PhyloCNN trained using 100K trees (paired Z-test; $p < 0.01$ ).
†Significantly higher absolute error compared to 2-Generation PhyloCNN trained using 10K trees (paired Z-test; $p < 0.01$ ).

MAE and MB values are shown for PhyloCNN using 100K and 10K training trees. We compared these results with those of DTL using CNN and CDV encoding (adapted from CBLV) and 1M, 100K, and 10K training trees. We used the same 1000 test trees for all methods, which we simulated according to the procedures described in the section *Simulating datasets for phylodynamics and macroevolution*.

The three options of PhyloCNN showed similar accuracies without significant differences when using the same number of training trees. PhyloCNN (2-Generation) trained with 100K trees significantly outperformed (p<0.01) all other methods, for all training set sizes, with respect to all parameters. When trained with 10K trees, PhyloCNN (2-Generation) outperformed (p<0.01) all other methods, for all training set sizes, with respect to all parameters except *q* (= *q*_01_=*q*_10_).

Again, the results show that the bias (in parentheses) induced by these different methods is small compared to the prediction errors.

**Table S7:** Results for the HIV dataset under the BDSS model.

|  | PhyloCNN<br>New HIV priors |  | PhyloDeep<br>Original priors |  | BEAST2<sup>1</sup> |
| --- | --- | --- | --- | --- | --- |
| Parameter | FFNN<br>SumStats | PhyloCNN | FFNN<br>SumStats<sup>1</sup> | CNN_CBLV<sup>1</sup> |  |
| $R_0$ | 1.98<br>[1.57-2.72] | 1.41<br>[1.16-1.74] | 1.60<br>[1.34–1.97] | 1.69<br>[1.40–2.08] | 1.41<br>[1.14-1.72] |
| $1/\gamma$ | 11.8<br>[8.9-15.5] | 11.3<br>[8.8-14.3] | 10.2<br>[8.3–12.8] | 9.8<br>[8.1–12.3] | 9.4<br>[7.6-11.7] |
| $X_{ss}$ | 15.8<br>[8.5-23.4] | 20.4<br>[10.6-26.2] | 8.8<br>[6.0–10.0] | 9.3<br>[6.7–10.0] | 14.5<br>[8.0-26.1] |
| $f_{ss}$ | 0.13<br>[0.08-0.17] | 0.13<br>[0.08-0.17] | 0.072<br>[0.05–0.12] | 0.079<br>[0.05–0.13] | 0.106<br>[0.05-0.17] |

<sup>1</sup>From (Voznica et al., 2022). See there for details on the use of BEAST2.

Point estimates and 95% confidence intervals (CI, in brackets) are shown for the basic reproductive number (*R*_0_), the infectious period (1/γ), the ratio of transmission rates between super- and normal-spreaders (*X*_SS_), and the fraction of super-spreaders (*f*_SS_). PhyloCNN (using the 2-Generation context) and FFNN_SumStats were trained using new priors and new simulated trees to account for the specificities of the HIV dataset (see text and Supp. Table S1 for explanations and details). The values presented here are the same as in Figure 6.

**Table S8:** Results for the primates dataset under the BiSSE model.

| Parameter | CNN_CDV<br>LL = -861.7 | MLE<br>LL = -846.9 | PhyloCNN<br>LL = -850.1 |
| --- | --- | --- | --- |
| $\lambda_0$ | 0.295 [0.222–0.394] | 0.395 [0.342–0.452] | 0.378 [0.310-0.440] |
| $\lambda_1$ | 0.093 [0.042–0.174] | 0.210 [0.167–0.266] | 0.166 [0.109-0.276] |
| $q$ | 0.009 [0.004–0.017] | 0.013 [0.008–0.018] | 0.011 [0.006-0.021] |
| $\tau$ | 0.234 [0.000–0.462] | 0.778 [0.690–0.858] | 0.610 [0.391-0.734] |

LL - Log-likelihood.

Point estimates and 95% confidence intervals (CI, in brackets) obtained with different methods are shown for turnover (τ), the speciation rates (λ_0_ and λ_1_, for states 0 and 1, respectively), and the symmetric class transition rate (*q*=*q*_01_=*q*_10_). PhyloCNN results were obtained using the 2-Generation context. Log-likelihood values corresponding to the point estimates of each method were calculated using diversitree 0.9-3 (Fitzjohn, 2012). The values presented here are the same as in Figure 7. See text for details.

---

**Figure S1:**
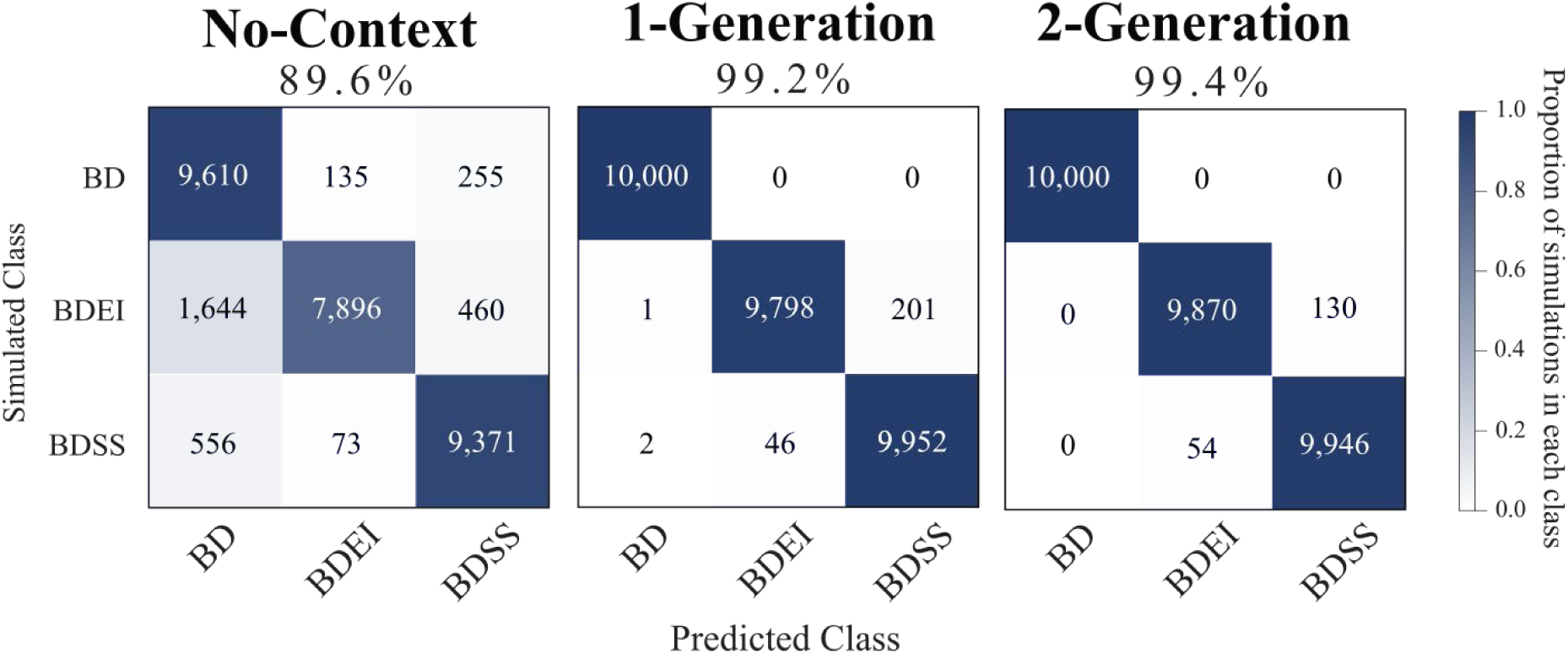
Selection of phylodynamics models with PhyloCNN. For model selection, PhyloCNN was trained using 10K training trees for each phylodynamics model (BD, BDEI, and BDSS). We observed a higher accuracy for both 1-Generation and 2-Generation than for No-Context. PhyloCNN (with both 1-Generation and 2-Generation) was more accurate than PhyloDeep trained using 4M trees per model (91.4% and 90.9% for FFNN_SumStats and CNN_CBLV, respectively; Voznica et al., 2022).

**Figure S2:**
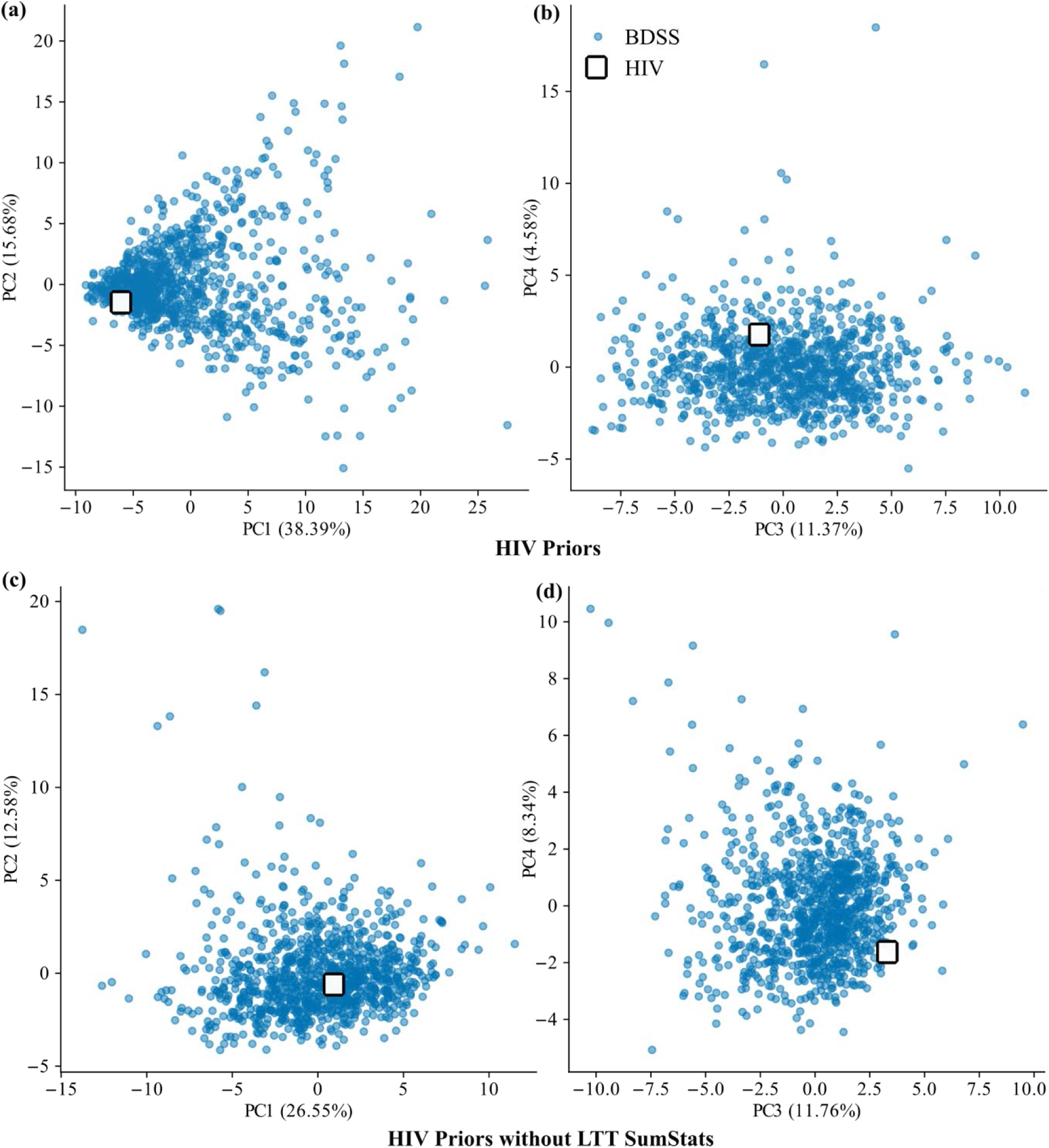
*A priori* model check for the HIV dataset. PCA analyses on the standardized SumStats obtained from 1K BDSS simulations under the new HIV priors (see text and Supp. Tab. S1 for details). Panels (a) and (b) include all 98 SumStats. Panels (c) and (d) do not include the SumStats corresponding to the lineages through time (LTT) plot. Panels (a) and (c): 1^st^ and 2^nd^ PCA components (PC1 and PC2). Panels (b) and (d): 3^rd^ and 4^th^ components (PC3 and PC4). The white square corresponds to the SumStats of the HIV dataset; it is included in the clouds for all panels. We performed PCA to check model adequacy without LTT SumStats because sampling in the HIV dataset is not random due to partner notification (Zhukova and Gascuel, 2024). In fact, we observed that 4 SumStats related to the y-axis of the LTT plot were rejected when we evaluated the 98 SumStats individually. For these 4 SumStats, the minimum and maximum values obtained from 10K BDSS simulations in the new priors do not bracket the value observed in the empirical HIV dataset. See https://github.com/manolofperez/phyloCNN for details.

**Figure S3:**
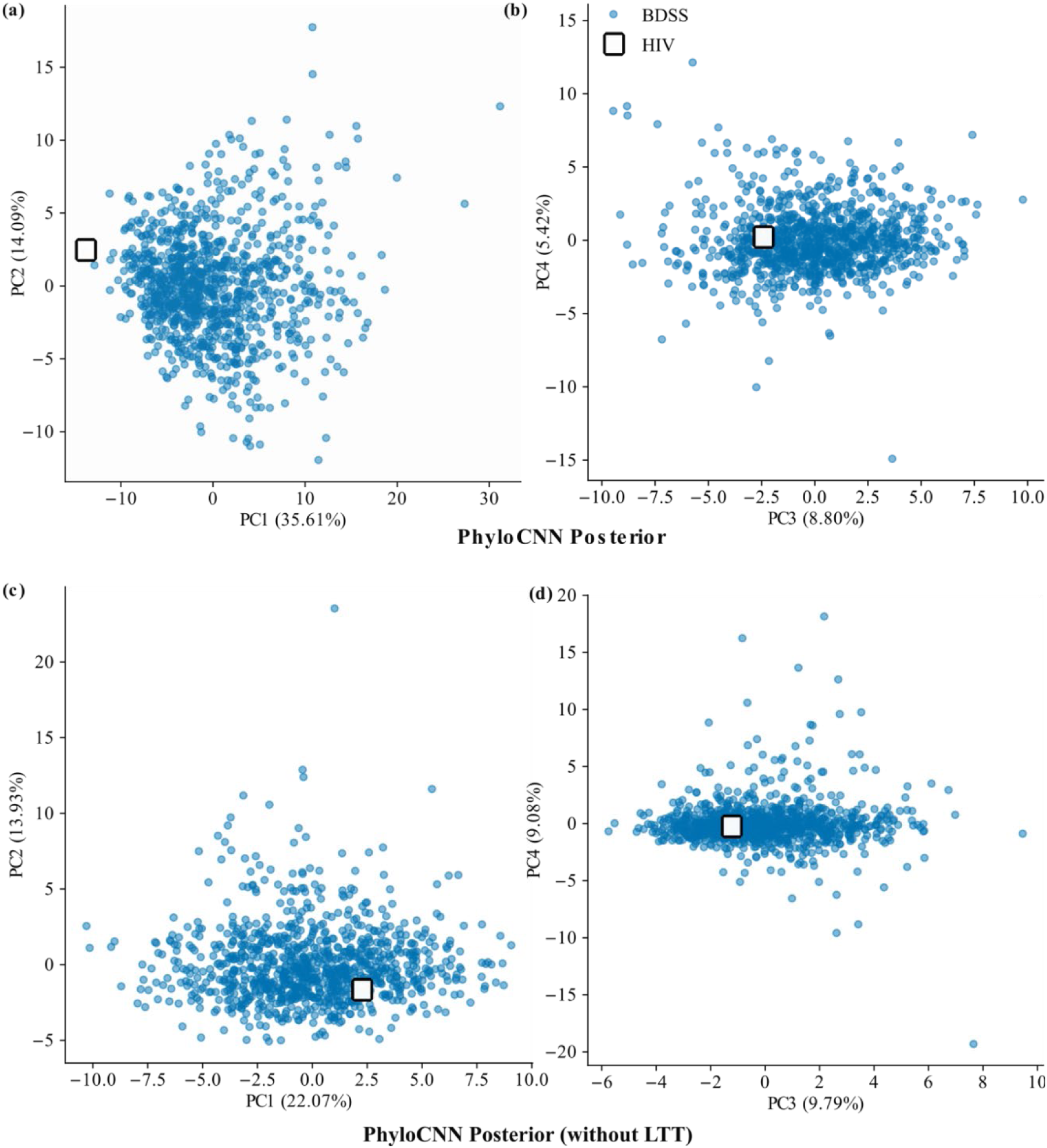
*A posteriori* model check for the HIV dataset. PCA analyses on the standardized SumStats from 1K BDSS simulations generated from the posterior distribution obtained with PhyloCNN for the HIV dataset (Figure 6). Panels (a) and (b) include all SumStats. Panels (c) and (d) do not include the SumStats corresponding to the lineages through time (LTT) plot. Panels (a) and (c): 1^st^ and 2^nd^ PCA components (PC1 and PC2). Panels (b) and (d): 3^rd^ and 4^th^ components (PC3 and PC4). The white square corresponds to the SumStats of the HIV dataset; it is included in the clouds for panels (b), (c) and (d), but somehow outside in (a) with respect to PC1 when including the LLT SumStats. Detailed analysis (https://github.com/manolofperez/phyloCNN) shows that 18 LTT SumStats are rejected; for these, the minimum and maximum values observed in the 1K simulations based on the posteriors do not bracket the observed value in the HIV dataset. This result was also observed in (Voznica et al., 2022) and is expected due to non-random sampling and partner notification in that dataset (Zhukova and Gascuel, 2024).

**Fig. S4.**
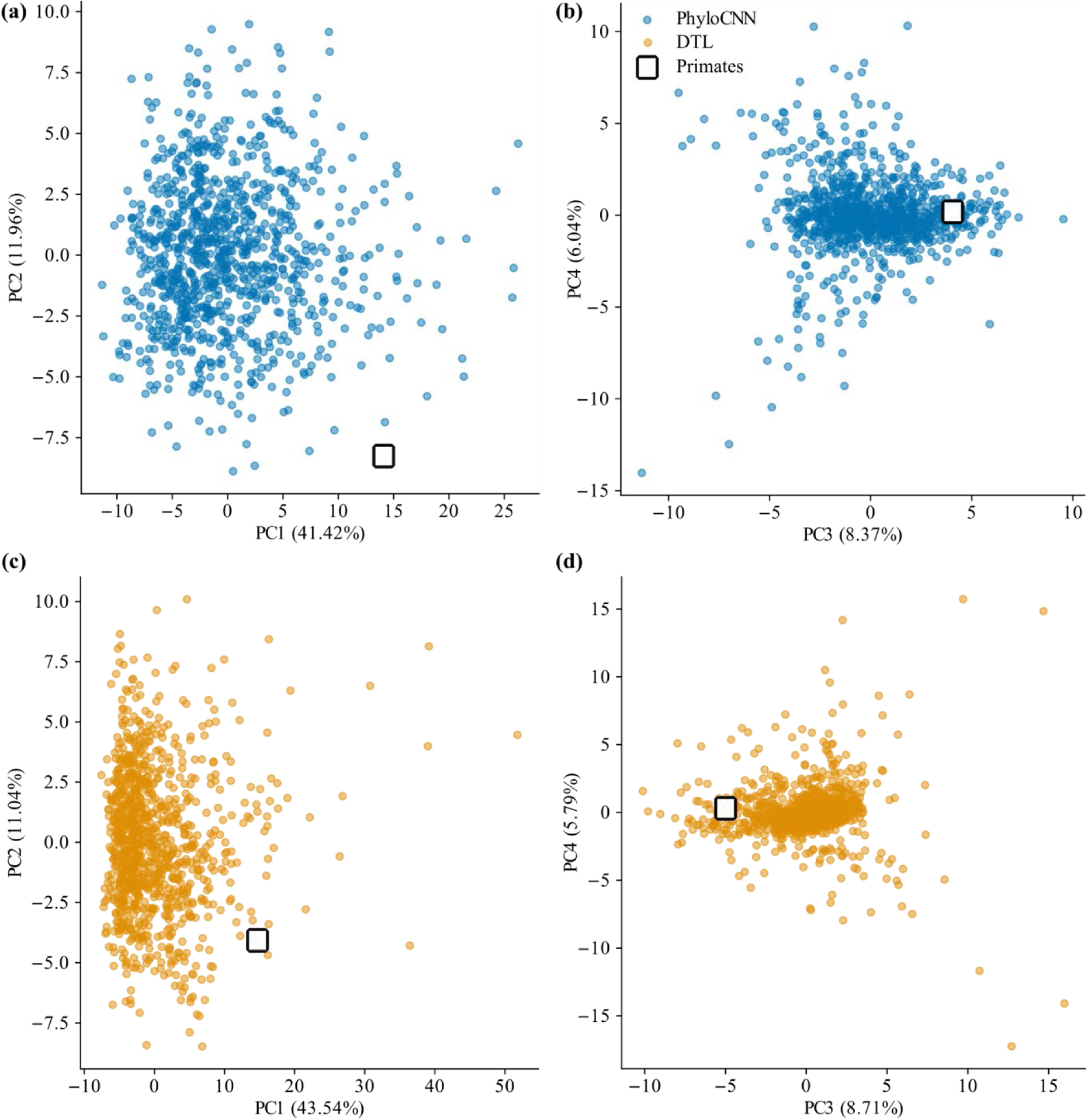
*A posteriori* model check for the primates dataset. PCA obtained from the 1K new simulations generated from the posterior distribution obtained with PhyloCNN (a, b) and DTL (c, d) for the primates dataset (Figure 7). Panels (a) and (c): 1^st^ and 2^nd^ PCA components (PC1 and PC2). Panels (b) and (d): 3^rd^ and 4^th^ components (PC3 and PC4). The white square corresponds to the SumStats of the primates dataset; although this primates point might appear outside the cloud of simulated points for PhyloCNN in (a) and DTL in (c), it is within the range of the simulations when we analyze each PC individually. Furthermore, when analyzed individually, all SumStats of the primates dataset are within the range of the SumStats from the posterior-based simulations (for both PhyloCNN and DTL).

## References

Attwood S.W., Hill-Cawthorne G., Khan A., Hayward A. 2022. Genomic epidemiology of SARS-CoV-2. Annu. Rev. Virol. 9:57–74.

Beaulieu J.M., O’Meara B.C. 2015. Extinction can be estimated from moderately sized molecular phylogenies. Evolution 69:1036–1043.

Beaumont M.A., Zhang W., Balding D.J. 2002. Approximate Bayesian computation in population genetics. Genetics 162:2025–2035.

Blum M.G.B. 2018. Regression approaches for ABC. In: Sisson S.A., Fan Y., Beaumont M.A., editors. Handbook of Approximate Bayesian Computation. Boca Raton (FL): Chapman and Hall/CRC; p.71–85.

Bouckaert R., Heled J., Kühnert D., Vaughan T., Wu C.H., Xie D., Suchard M.A., Rambaut A., Drummond A.J. 2014. BEAST 2: a software platform for Bayesian evolutionary analysis. PLoS Comput. Biol. 10.

Chollet F. 2017. Xception: deep learning with depthwise separable convolutions. In: Proceedings of the IEEE Conference on Computer Vision and Pattern Recognition. p.1251–1258.

Clevert D.A., Unterthiner T., Hochreiter S. 2015. Fast and accurate deep network learning by exponential linear units (ELUs). arXiv 1511.07289.

Condamine F.L., Rolland J., Morlon H. 2019. Assessing the causes of diversification slowdowns: temperature-dependent and diversity-dependent models receive equivalent support. Ecol. Lett. 22:1900–1912.

Csilléry K., Blum M.G.B., Gaggiotti O.E., François O. 2010. Approximate Bayesian computation (ABC) in practice. Trends Ecol. Evol. 25:410–418.

Csilléry K., François O., Blum M.G.B. 2012. abc: an R package for approximate Bayesian computation (ABC). Methods Ecol. Evol. 3:475–479.

Danesh G., Saulnier E., Gascuel O., Choisy M., Alizon S. 2023. TiPS: rapidly simulating trajectories and phylogenies from compartmental models. Methods Ecol. Evol. 14:487–495.

De Maio N., Wu C.H., O’Reilly K.M., Wilson D.J. 2015. New routes to phylogeography: a Bayesian structured coalescent approximation. PLoS Genet. 11.

Etienne R.S., Haegeman B., Stadler T., Aze T., Pearson P.N., Purvis A., Phillimore A.B. 2012. Diversity-dependence brings molecular phylogenies closer to agreement with the fossil record. Proc. R. Soc. B 279:1300–1309.

Fabre P.-H., Rodrigues A., Douzery E.J.P. 2009. Patterns of macroevolution among primates inferred from a supermatrix of mitochondrial and nuclear DNA. Mol. Phylogenet. Evol. 53:808–825.

Featherstone L.A., Zhang J.M., Vaughan T.G., Duchêne S. 2022. Epidemiological inference from pathogen genomes: a review of phylodynamic models and applications. Virus Evol. 8.

Featherstone L.A., Duchene S., Vaughan T.G. 2023. Decoding the fundamental drivers of phylodynamic inference. Mol. Biol. Evol. 40:msad132.

Fitzjohn R. G. 2012. Diversitree: comparative phylogenetic analyses of diversification in R. Methods in Ecology and Evolution, 3:1084–1092.

Flagel L., Brandvain Y., Schrider D.R. 2019. The unreasonable effectiveness of convolutional neural networks in population genetic inference. Mol. Biol. Evol. 36:220–238.

Garrick R.C., Bonatelli I.A.S., Hyseni C., Morales A., Pelletier T.A., Perez M.F., Rice E., Satler J.D., Symula R.E., Thomé M.T.C., Carstens B.C. 2015. The evolution of phylogeographic data sets. Mol. Ecol. 24:1164–1171.

Gascuel O., Steel M. 2014. Predicting the Ancestral Character Changes in a Tree is Typically Easier than Predicting the Root State. Syst. Biol. 63:421–435.

Gómez J.M., Verdú M. 2012. Mutualism with plants drives primate diversification. Syst. Biol. 61:567–577.

Grenfell B.T., Pybus O.G., Gog J.R., Wood J.L.N., Daly J.M., Mumford J.A., Holmes E.C. 2004. Unifying the epidemiological and evolutionary dynamics of pathogens. Science 303:327–332.

Haykin S. 1999. Neural Networks: A Comprehensive Foundation. 2nd ed. Upper Saddle River (NJ): Prentice Hall.

Höhna S., Stadler T., Ronquist F., Britton T. 2011. Inferring speciation and extinction rates under different sampling schemes. Mol. Biol. Evol. 28:2577–2589.

Huerta-Cepas J., Serra F., Bork P. 2016. ETE 3: reconstruction, analysis, and visualization of phylogenomic data. Mol. Biol. Evol. 33:1635–1638.

Janzen T., Etienne R.S. 2024. Phylogenetic tree statistics: A systematic overview using the new R package ‘treestats’. Mol. Phylogenet Evol. 200:108168.

Kingma D.P., Ba J. 2014. Adam: a method for stochastic optimization. arXiv:1412.6980.

Kirschner P., Perez M.F., Záveská E., Sanmartín I., Marquer L., Schlick-Steiner B.C., Alvarez N., STEPPE Consortium, Steiner F.M., Schönswetter P. 2022. Congruent evolutionary responses of European steppe biota to late Quaternary climate change. Nat. Commun. 13: 1921.

Korfmann K., Sellinger T.P.P., Freund F., Fumagalli M., Tellier A. 2024. Simultaneous inference of past demography and selection from the ancestral recombination graph under the Beta coalescent. Peer Community Journal 4:e33.

Lajaaiti I., Lambert S., Voznica J., Morlon H., Hartig F. 2023. A Comparison of Deep Learning Architectures for Inferring Parameters of Diversification Models from Extant Phylogenies. bioRxiv 2023.03.03.530992.

Lambert S., Voznica J., Morlon H. 2023. Deep learning from phylogenies for diversification analyses. Syst. Biol. 72:1262–1279.

Landis M., Thompson A. 2024. phyddle: software for phylogenetic model exploration with deep learning. BioRxiv 10.1101/2024.08.06.606717.

Louca S., Doebeli M. 2018. Efficient comparative phylogenetics on large trees. Bioinformatics 34:1053–1055.

Louca S., Pennell M.W. 2020a. Extant timetrees are consistent with a myriad of diversification histories. Nature 580:502–505.

Louca S., Pennell M.W. 2020b. Why extinction rates should be estimated from molecular phylogenies. Proc. Natl. Acad. Sci. U.S.A. 117:24194–24200.

MacPherson A., Louca S., McLaughlin A., Joy J.B., Pennell M.W. 2022. Unifying phylogenetic birth-death models in epidemiology and macroevolution. Syst. Biol. 71:172–189.

Maddison W.P., Midford P.E., Otto S.P. 2007. Estimating a binary character’s effect on speciation and extinction. Syst. Biol. 56:701–710.

Mo Z., Siepel A. 2023. Domain-adaptive neural networks improve supervised machine learning based on simulated population genetic data. PLoS Genet 19(11): e1011032.

Mo Y.K., Hahn M.W., Smith M.L. 2024. Applications of machine learning in phylogenetics. Mol. Phylogenet Evol. 196:108066.

Mondal M., Bertranpetit J., Lao O. 2019. Approximate Bayesian computation with deep learning supports a third archaic introgression in Asia and Oceania. Nat. Commun. 10:246.

Morlon H, Potts MD, Plotkin JB. 2010. Inferring the dynamics of diversification: a coalescent approach. PLoS Biol. 8:e1000493.

Morlon H. 2014. Phylogenetic approaches for studying diversification. Ecol. Lett. 17:508– 525.

Morlon H., Andréoletti J., Barido-Sottani J., Lambert S., Perez-Lamarque B., Quintero I., Senderov V., Véron P. 2024. Phylogenetic insights into diversification. Annu. Rev. Ecol. Evol. Syst. 55.

Müller N.F., Rasmussen D.A., Stadler T. 2017. The structured coalescent and its approximations. Mol. Biol. Evol. 34:2970–2981.

Nee S. 2006. Birth-death models in macroevolution. Annu. Rev. Ecol. Evol. Syst. 37:1–17.

Oliveira E.A., Perez M.F., Bertollo L.A.C., Gestich C.C., Ráb P., Ezaz T., Souza F.H.S., Viana P., Feldberg E., Oliveira E.H.C., Cioffi M.B. 2020. Historical demography and climate-driven distributional changes in a widespread Neotropical freshwater species with high economic importance. Ecography 43:1291–1304.

Pennell M.W., Harmon L.J. 2013. An integrative view of phylogenetic comparative methods: connections to population genetics, community ecology, and paleobiology. Ann. N.Y. Acad. Sci. 1289:90–105.

Perez M.F., Bonatelli I.A.S., Romeiro-Brito M., Franco F.F., Taylor N.P., Zappi D.C., Moraes E.M. 2022. Coalescent-based species delimitation meets deep learning: insights from a highly fragmented cactus system. Mol. Ecol. Resour. 22:1016–1028.

Prangle D., Everitt R.G., Kypraios T. 2018. A rare event approach to high-dimensional approximate Bayesian computation. Stat. Comput. 28:819–834.

Pudlo P., Marin J.M., Estoup A., Cornuet J.M., Gautier M., Robert C.P. 2016. Reliable ABC model choice via random forests. Bioinformatics 32:859–866.

Rabosky D.L. 2014. Automatic detection of key innovations, rate shifts, and diversity-dependence on phylogenetic trees. PloS One. 9:e89543.

Rasmussen D.A., Kouyos R., Günthard H.F., Stadler T. 2017. Phylodynamics on local sexual contact networks. PLoS Comput. Biol. 13.

Robert C.P., Cornuet J.M., Marin J.M., Pillai N.S. 2011. Lack of confidence in approximate Bayesian computation model choice. Proc. Natl. Acad. Sci. U.S.A. 108:15112– 15117.

Sanchez T., Cury J., Charpiat G., Jay F. 2021. Deep learning for population size history inference: design, comparison, and combination with approximate Bayesian computation. Mol. Ecol. Resour. 21:2645–2660.

Sanmartín I, Meseguer AS. 2016. Extinction in phylogenetics and biogeography: from timetrees to patterns of biotic assemblage. Front. Genet. 7:35.

Saulnier E., Gascuel O., Alizon S. 2017. Inferring epidemiological parameters from phylogenies using regression-ABC: a comparative study. PLoS Comput. Biol. 13.

Schrider D.R., Kern A.D. 2018. Supervised machine learning for population genetics: a new paradigm. Trends Genet. 34:301–312.

Scire J., Barido-Sottani J., Kühnert D., Vaughan T.G., Stadler T. 2022. Robust phylodynamic analysis of genetic sequencing data from structured populations. Viruses 14:1648.

Sheehan S., Song Y.S. 2016. Deep learning for population genetic inference. PLoS Comput. Biol. 12.

Smith M.L., Hahn M.W. 2023. Phylogenetic inference using generative adversarial networks. Bioinformatics 39:btad543

Stadler T. 2010. Sampling-through-time in birth-death trees. J. Theor. Biol. 267:396–404.

Stadler T., Kühnert D., Bonhoeffer S., Drummond A. J. 2012. Birth–death skyline plot reveals temporal changes of epidemic spread in HIV and hepatitis C virus (HCV). Proc. Natl. Acad. Sci. U.S.A. 110:228–233.

Stadler T., Bonhoeffer S. 2013. Uncovering epidemiological dynamics in heterogeneous host populations using phylogenetic methods. Philos. Trans. R. Soc. B 368:20120198.

Stadler T., Kühnert D., Rasmussen D.A., du Plessis L. 2014. Insights into the early epidemic spread of Ebola in Sierra Leone provided by viral sequence data. PLoS Curr 6.

Stadler T., Bokma F. 2013. Estimating Speciation and Extinction Rates for Phylogenies of Higher Taxa. Syst. Biol. 62: 220–230.

Stadler T., Kühnert D., Rasmussen D.A., du Plessis L. 2014. Insights into the early epidemic spread of Ebola in Sierra Leone provided by viral sequence data. PLoS Curr. 6.

Swiss HIV Cohort Study; Schoeni-Affolter F, Ledergerber B, Rickenbach M, Rudin C, Günthard HF, Telenti A, Furrer H, Yerly S, Francioli P. (2010). Cohort profile: the Swiss HIV Cohort study. Int J Epidemiol. 39:1179–89.

Thompson A., Liebeskind B.J., Scully E.J., Landis M.J. 2024. Deep learning and likelihood approaches for viral phylogeography converge on the same answers whether the inference model is right or wrong. Syst. Biol. 73:183–206.

Torada L., Lorenzon L., Beddis A., Isildak U., Pattini L., Mathieson S., Fumagalli M. 2019. ImaGene: a convolutional neural network to quantify natural selection from genomic data. BMC Bioinformatics 20:337.

Vaughan T.G., Scire J., Nadeau S.A., Stadler T. 2024. Estimates of early outbreak-specific SARS-CoV-2 epidemiological parameters from genomic data. Proceedings of the National Academy of Sciences. 121:e2308125121.

Volz E.M., Koelle K., Bedford T. 2013. Viral phylodynamics. PLoS Comput. Biol. 9.

Voznica J., Zhukova A., Boskova V., Saulnier E., Lemoine F., Moslonka-Lefebvre M., Gascuel O. 2022. Deep learning from phylogenies to uncover the epidemiological dynamics of outbreaks. Nat. Commun. 13:3896.

Xie R., Adam D.C., Hu S., Cowling B.J., Gascuel O., Zhukova A., Dhanasekaran V. 2024. Integrating contact tracing data to enhance outbreak phylodynamic inference: a deep learning approach. Mol. Biol. Evol. 10.1093/molbev/msae232 (Epub ahead of print).

Ying Z., You J., Morris C., Ren X., Hamilton W., Leskovec J. 2018. Hierarchical Graph Representation Learning with Differentiable Pooling. arXiv 1806.08804.

Zhukova A., Hecht F., Maday Y., Gascuel O. 2023. Fast and accurate maximum-likelihood estimation of multi-type birth–death epidemiological models from phylogenetic trees. Syst. Biol. 72:1387–1402.

Zhukova A., Gascuel O. 2024. Accounting for partner notification in epidemiological birth-death models. medRxiv 2024.09.09.24313296.

